# Machine Learning Uncovers Vascular Endothelial Cell Identity Genes by Expression Regulation Features in Single Cells

**DOI:** 10.1101/2024.08.27.609808

**Authors:** Kulandaisamy Arulsamy, Bo Xia, Hong Chen, Lili Zhang, Kaifu Chen

## Abstract

Deciphering cell identity genes is pivotal to understanding cell differentiation, development, and many diseases involving cell identity dysregulation. Here, we introduce SCIG, a machine-learning method to uncover cell identity genes in single cells. In alignment with recent reports that cell identity genes are regulated with unique epigenetic signatures, we found cell identity genes exhibit distinctive genetic sequence signatures, e.g., unique enrichment patterns of cis-regulatory elements. Using these genetic sequence signatures, along with gene expression information from single-cell RNA-seq data, enables SCIG to uncover the identity genes of a cell without a need for comparison to other cells. Cell identity gene score defined by SCIG surpassed expression value in network analysis to uncover master transcription factors regulating cell identity. Applying SCIG to the human endothelial cell atlas revealed that the tissue microenvironment is a critical supplement to master transcription factors for cell identity refinement. SCIG is publicly available at https://github.com/kaifuchenlab/SCIG, offering a valuable tool for advancing cell differentiation, development, and regenerative medicine research.

## Highlights

- Cell identity genes exhibit unique genetic sequence codes for their cis-regulations.
- Machine learning accurately uncovers cell identity genes by their cis-regulatory signatures.
- Cell identity gene score surpassed expression value in network analysis to uncover master transcription factors regulating cell identity.
- Landscape analysis of cell identity genes revealed insight into cell identity refinement by tissue microenvironment as a supplement to master transcription factors.

## Introduction

Every cell type possesses a distinct collection of several hundred or more cell identity genes (CIGs) governing its cellular characteristics. A cell might express both its CIGs and many other gene categories such as housekeeping genes and heat shock genes. Accurate expression of CIGs is essential for steering cell differentiation and the development of tissues and organs in living organisms. The CIGs of a cell comprise an intricate gene regulatory network (GRN), where several master transcription factors coordinate the expression of the CIGs(1–3). For instance, the expression program of pluripotent stem cells can be established by the master transcription factors Oct4, Sox2, Klf4, and c-Myc(4, 5). Cell differentiation from stem cells to somatic cell types, trans-differentiation between somatic cell types, and reprogramming from somatic cell types to stem cells, are orchestrated by their CIG regulatory networks, enabling a source cell type to give rise to a specific target cell type(4). This dynamic process encompasses silencing and activating the CIGs of the source and target cell types, respectively. Uncovering the CIGs of individual cell types is pivotal in regenerative medicine to facilitate the precise generation of desired target cell types from a source cell type(6, 7).

It is increasingly recognized that the same type of epigenetic modification tends to display different patterns between CIGs and most of the other expressed genes in the same cell. For instance, the histone modification H3K4me3 tends to display a broad enrichment pattern of about 5kb to 100kb covering both the promoter and the gene body of a CIG, whereas H3K4me3 tends to display a sharp enrichment in only about 1kb promoter region at each of the other expressed genes(8, 9). It is also reported that CIGs tend to be regulated by super-enhancers (5, 10), which show a broad enrichment pattern of the histone modification H3K4me1 and H3K27ac covering a long stretch of enhancers, while most other genes are regulated by typical enhancers each displaying narrow enrichment peaks of these histone modifications(8, 9). Moreover, CIGs tend to show the lowest RNA stability in the cell because the m6A methyltransferase preferentially modifies these RNAs in a co-transcriptional manner, likely guided by chromatin features, such as the super-enhancers and broad H3k4me3 modification(11). Researchers have been able to uncover CIGs in a cell type using these epigenetic profiles(12). However, generating these epigenetic profiles often entails a substantial investment of time and resources. Moreover, profiling multiple histone modifications in parallel from each single cell at the genome-wide scale is still a technological challenge. Therefore, uncovering cell identity genes by their epigenetic profiles, especially when applied to single-cell study, may be limited due to the difficulty of generating the epigenetic profile data.

Uncovering CIGs solely based on their expression in a cell also presents a great challenge because the expression level might resemble those of many other genes such as the housekeeping genes. Existing computation solutions often aim at defining cell type-specific marker genes by comparison between cell types at the bulk level or between cell clusters in a single-cell RNA sequencing (scRNA-seq) dataset. These solutions are not optimal because different CIGs display great differences in the degree of expression specificity. For instance, c-Myc is a well-known identity gene of embryonic stem cells(4), but also expressed to regulate the identity or play functions in some other cell types, such as the Fibroblast (13, 14) and hematopoietic stem cell (15, 16). Cell identity is often governed by the cell type-specific combination of identity genes, for which the combination is cell type-specific, but many CIGs might be not strictly cell type-specific. Therefore, some CIGs might be missed from the list of cell type-specific genes.

Furthermore, the defined cell type-specific genes for a cell type may vary when compared to different sets of other cell types. It is also a challenge to distinguish between cell type-specific and experimental condition-specific genes, e.g., heat shock genes. Other bioinformatics methods aim at constructing the gene expression regulation network to define pivotal regulators^3^. This solution is yet not optimal for defining CIGs, because the gene regulation network may include both a sub-network of CIGs and some sub-networks of other gene categories, e.g., cell cycle genes, stress response genes, and genes for DNA damage repair. It is yet a challenge to distinguish between these sub-networks. Therefore, despite the widespread application of scRNA-seq in recent years to define cell type-specific genes or expression networks, there is no existing method that employs a single-cell transcriptome to directly uncover CIGs.

Genes belonging to some functional categories were reported to show some unique genetic sequence features. For example, housekeeping genes tend to show lower promoter sequence conservation(17), sequences with less potential for nucleosome formation in the promoter region(18), and shorter gene bodies(19–21). Recent studies demonstrated that genetic sequence signatures hold promise to be cis-regulatory codes for some aspects of biological function, e.g., RNA structure(22–24), protein-binding(25), RNA stability(26), and promoters and enhancers activities(27, 28). Intriguingly, CIG tends to show a broad enrichment of transcription factor binding motifs across the gene body, in contrast to the narrow enrichment of these motifs in only the promoters of most other genes(8). Exploring these genetic sequence signatures has provided valuable clues to understanding CIG expression regulation and might facilitate the identification of CIGs. Therefore, we reasoned that integrating gene expression information with genetic sequence codes to uncover CIGs may be a promising approach offering significant advantages over prior methods.

In this study, we demonstrated the effectiveness of combining genetic sequence signatures with expression information for distinguishing between CIGs and the rest of the genes. Building upon this characterization, we developed a robust machine-learning algorithm, SCIG, for uncovering Single-Cell Identity Genes. Applying SCIG to diverse single-cell datasets, including individual subtypes of the endothelial cell lineage, our analysis generated new insights into the expression regulation of CIGs in cell differentiation. Notably, cell clustering results based on CIG scores more accurately reflected cell types and their differentiation trajectory when compared to the conventional strategy based on RNA expression values alone. The CIG catalog uncovered by SCIG holds great promise for advancing cell identity reprogramming experiments and facilitating the generation of desired cell types in regenerative medicine.

## Materials and Methods

### Analyzing the Genetic and Expression Features of Cell Identity Genes

We obtained a set of CIGs and control genes manually curated for 10 human cell types in a recent work(12) (**Supplementary Table 1 and 2**). We computed five categories of features from RNA expression and gene sequence data to capture the characteristics of CIGs.

**i. Gene expression values:** The bulk RNA-seq raw reads for 10 human cell types, including H1-hESC, CD34+ HPC, B cell, HUVECs, human mammary epithelial cells, neural cells, MRG cell, normal human lung fibroblast, mesenchymal stem cell, and human skeletal muscle myoblast, were collected from the Encyclopedia of DNA Elements (ENCODE) project(29) and NCBI Sequence Read Archive (SRA)(30) repositories **(Supplementary Table3)**. The human reference genome version hg38.104 was downloaded from the Ensembl database(31), and the longest transcript for each protein-coding gene was selected for further analysis. We employed Trim Galore (version 0.6.6, https://www.bioinformatics.babraham.ac.uk/projects/trim_galore/) for adaptor trimming and removal of low-quality sequencing reads, followed by mapping the high-quality reads to the human reference genome using the STAR alignment tool (version 2.7.9a)(32). Quantification of the mapped reads was performed using FeatureCounts (v2.0.3)(33) for the entire gene and exonic regions. The quantified read counts were then converted into log2-transformed Transcript per million (TPM) values for downstream analyses. For the mouse genome, expression data for the aforementioned 10 cell types was gathered from the Mouse Cell Atlas (MCA) database (34).
**ii. Gene expression specificity metrics**: Gene expression values for human tissues and cell types were sourced from various RNA-seq projects, including the Genotype-Tissue Expression (GTEx)(35), Functional Annotation of Mammalian Genomes 5 (FANTOM5)(36), and Human Protein Atlas(37, 38). These data were downloaded from The Human Protein Atlas website (https://www.proteinatlas.org/about/download) (37, 38). For mouse tissues and cell types, the expression data was obtained from MCA(34), Mouse ENCODE(39), and mammalian transcriptomic database(40). Next, the tspex(41) python package was used to calculate various gene expression specificity metrics, including tau, gini coefficient, simpson index, shannon specificity, and roku specificity, yielding specificity scores ranging from 0 to 1. Higher scores indicate that genes are expressed more specifically in certain tissues or cell types.
**iii. Binding motifs and sites of transcription factors, RNA-binding proteins, and miRNAs:** The human and mouse transcription factor binding motifs were obtained from the “MotifDb” (42) package and were used to scan the entire human and mouse genome sequences. The analysis focused on determining the number of transcription factor binding motifs within different genomic regions, including the promoter region with various window lengths (500bp to 5kb in both upstream and downstream sequences), gene bodies, 5’-UTR, exons, introns, and 3’-UTR. The binding sites of RNA-binding proteins were downloaded from the oRNAment database(43), and the number of binding sites within specific genomic regions (5’-UTR, exons, introns, and 3’-UTR) was determined using the bedtools intersect command(44). Additionally, miRWalk(45) was employed to retrieve miRNA binding sites. Thereafter, the number of miRNA binding sites within the 5’-UTR, coding sequence (CDS), and 3’-UTR regions was calculated. The number of these features were normalized based on the length of the respective genomic regions.
**iv. Evolutionary features:** The phyloP100way sequence conservation score for the human and mouse genome was obtained from the UCSC database(46). Using in-house scripts, we computed the mean and median conservation scores for a whole gene sequence and different genomic regions.
**v. Generic gene-level features:** The gene architecture features, including gene sequence length and the lengths of different genomic elements (exon, intron, CDS, 5’-UTR, and 3’-UTR), were calculated from the human and mouse reference genome(31). Additionally, the number of introns and exons, as well as the ratio of introns to exons, were also computed. For each gene, we computed the frequencies of single-, di-, and tri-nucleotides in the gene sequence. Codon biases were determined by grouping triplet codons based on their coding amino acid and calculating their total proportion, mean, median, and coefficient of variation. Additionally, we included the transcription start site (TSS) distance, which represents the number of base pairs between the TSS of a gene and its closest TSS in the chromatin.

### Features Filtering and Selection

Based on the feature extraction procedures described above, a total of 680 features were obtained for each gene. Initially, a Wilcoxon(47) nonparametric test was conducted to determine the significance of the difference in each feature between CIGs versus control or housekeeping genes. Features with a P-value greater than 0.05 when comparing CIGs to control or housekeeping genes were subsequently removed. Next, we calculated the Pearson correlation coefficient between features. For a pair of features with a correlation coefficient above 0.90, the feature with the greater P-value of its difference between CIGs and control genes was removed. These steps resulted in a final set of 73 non-redundant and significant features.

In our pursuit of identifying potential features for machine learning model development, we employed a systematic search followed by a forward feature selection algorithm(48). The systematic search involved exploring all possible combinations of three features, resulting in a selection of the top 10, 000 combinations based on the performance metric, Matthew’s correlation coefficient, in 10-fold cross-validation. Subsequently, we implemented the forward selection algorithm to gradually incorporate additional features from the pool into the selected combinations. The forward feature selection process continued until no further enhancement in model performance was observed, resulting in a final set of selected features. To assess the efficacy of this feature selection method, we compared it with several other approaches, including the SK-best-mutual information, ANOVA F-classification, forward sequential selection (FSS), backward stepwise elimination (BSE), recursive feature elimination based on Logistic Regression (RFE LR) with L1 regularization(49).

### Developing the SCIG, a Logistic Regression Model to Uncover CIGs

After feature selection, the training dataset was standardized using Sklearn’s preprocessing libraries(49). A logistic regression algorithm with an L1 penalty parameter was applied to develop the model, aiming to prevent overfitting. The model selection involved a 10-fold cross-validation procedure, and further validation was conducted through bootstrapping with 500 iterations, using an 80% training and 20% testing split. Model performance was assessed using statistical metrics such as sensitivity, specificity, accuracy, Matthew’s correlation coefficient, F1-score, and area under the receiver operating characteristic curve (AUROC).

### Collecting Reported Master Transcription Factors of Cell Identity

The human transcription factors were obtained from the humantfs database (http://humantfs.ccbr.utoronto.ca/) (50). We extracted the reported master transcription factors of cell identity manually curated for 10 cell types in recent work(12). The remaining transcription factors from the human transcription factor list were recognized as the control transcription factors in these 10 cell types. As the number of control transcription factors exceeds the number of master transcription factors, we addressed the class imbalance issue by utilizing the SMOTE algorithm(51), resulting in a balanced dataset of 221 randomly selected master and control transcription factors for model development.

### Machine Learning Model to Uncover Master Transcription Factors in CIG Networks

We compiled a reference gene regulatory network (GRN) from seven different resources, including RegNetwork(52), DoRothEA(53), CellNet(3), GRNdb(54), ANANSE(55), PANDA(56), and Huang, J.K. et al(57). We focused on protein-coding gene interactions and determined the reliability of each interaction based on the number of votes from these sources, taking only the interactions that were present in at least two sources. For each known master transcription factors and control transcription factor, we extracted their corresponding GRNs from the compiled reference GRN. Next, we computed several features including the number of children edges (indicating the number of target genes regulated by a specific transcription factor), the number of parent edges (indicating the number of genes regulating a specific transcription factor), cell identity gene score (CIG score) from SCIG, and RNA expression of the genes. Furthermore, we calculated the mean, median, and coefficient of variation for each feature of the transcription factors. This process yielded a total of 23 features for each master and control transcription factor, which were subsequently used for constructing a logistic regression model to learn network features enriched in master transcription factors. Important features to use in the model were selected through systematic feature selection using 10-fold cross-validation. The model’s performance was evaluated using statistical measures like accuracy, Matthews correlation coefficient, and F1-score during both 10-fold cross-validation and bootstrapping procedures.

### Exploring Single-Cell Identity Gene Networks in Human Fetal Heart Using hdWGCNA

Firstly, we employed the SCIG algorithm to calculate cell identity scores for individual genes in individual cells from human fetal hearts(58) (**Figure 4a**). Then, we supplied the CIG score matrix and expression matrix independently to hdWGCNA(59) for gene network analysis at the single-cell level. Initially, we selected the genes that expressed in a minimum of 5% of cells in both the expression and CIGs score matrices. Subsequently, we generated metacells using the ‘MetacellsByGroups (k=20, min_cells=30)’ function, followed by normalization using ‘NormalizeMetacells’. Notably, normalization was omitted for the CIG score matrix as it was inherently in normalized form. Further, we constructed the network by considering all cells with default options. Each identified network module was visualized using the ‘ModuleNetworkPlot’ function.

### Using Cell Identity Gene Score for Single-Cell Clustering Analysis

The human forebrain glutamatergic neurogenesis(60) dataset consisted of 1720 scRNA-seq profiles, which were utilized to predict CIGs using the SCIG algorithm. This transformed the gene-cell expression matrix into a gene-cell CIG score matrix. We then analyzed the expression matrix and CIG score matrix independently using the Seurat(61) package for cell clustering and cell type annotation. Only protein-coding genes were analyzed, while mitochondrial and ribosomal genes were filtered out. For the expression matrix, expression values were log-normalized, and the top 2000 highly variable genes were selected using the “NormalizeData” and “FindVariableFeatures” functions, respectively, with default parameters. The normalized expression matrix was then scaled and centered using the “ScaleData” function, followed by running the “RunPCA (reduction = “pca”, dims = 1:40)”, “RunUMAP”, “FindNeighbors”, and “FindClusters” function to obtain cell clusters. In the case of the CIG score matrix, the same clustering pipeline was employed except that the log-normalization step was skipped, as the CIG scores were already standardized. The identified cell clusters were annotated based on their RNA expression levels or CIG scores. The cell types identified based on each matrix were further analyzed using ScVelo(62) to determine their future state or transition direction based on spliced and unspliced mRNA expression values. The velocity information obtained was then utilized in CellRank(63) to identify the potential initial and terminal sites/cells within the given cell types. Additionally, CellRank quantified the transition probability for each terminal site from all other cell types, providing valuable insights into cell dynamics and potential cell state transitions.

### CIG Analysis in the Process of Endothelial Differentiation

We obtained scRNA-seq profiles during endothelial differentiation from human embryonic stem cells (H9) at multiple time points, including days 0 (embryonic stem cells), 4 (mesoderm), 6 (mesenchymal), 8 (endothelial cell progenitors), and 12 (endothelial cells) (64). The unique molecular identifiers (UMI) count matrix were preprocessed and transformed into pseudo-bulk datasets for each time point. These datasets were subsequently utilized in the SCIG model to predict CIGs and their master transcription factors.

### Endothelial Cell Identity Gene Landscape Analysis in 15 Human Tissue Types

We obtained scRNA-seq data from 15 tissues, encompassing adipose, bladder, breast, gut, heart, intestine, kidney, liver, lung, ovary, skeletal muscle, skin, stomach, testis, and thymus, sourced from the DISCO(65) database. Endothelial subtypes, including arterial, capillary, venous, and lymphatic, were specifically extracted from each tissue. Subsequently, we aggregated gene UMI counts for each endothelial cell subtype from the count matrix. This consolidated dataset was then integrated into SCIG for the prediction of CIGs and their master transcription factors. To determine the specificity of each cell identity gene and their master transcription factors across 15 tissues, we retrieved the SCIG-derived scores for the genes (with false discovery rate (FDR)<0.05) in each endothelial subtype. Subsequently, these retrieved scores of cell identity genes and master transcription factors were utilized for Tau score calculation using tspex(41). We conducted the Wilcoxon nonparametric one-sided test for assessing the statistical difference between the Tau specificity score of CIGs and CIG master transcription factors.

### Pathway Enrichment Analysis

The significant CIGs (with FDR<0.05), high-expression genes, and highly variable genes were subjected to a pathway enrichment analysis using the ClusterProfiler(66) R package. The enriched Gene Ontology (GO) pathways were determined based on the least adjusted P-values using the Benjamini method, with a significance threshold of q-value <0.05. This pipeline helps identify the biological processes and functions associated with the identified CIGs and other genes. The fold enrichment for each pathway was computed by calculating the ratio between the gene ratio and the background ratio obtained from ClusterProfiler output.

### Statistical Analysis

For each genetic sequence and expression-derived feature, we used two-tailed Wilcoxon’s test(47) for assessing the statistical difference (P-value) between CIGs and housekeeping, control genes. From SICG output, the significant CIGs and their master transcription factors were identified by FDR <0.05. We utilized pROC(67) package for determining the P-values of ROC curves. During the pathway enrichment analysis, we used the q-value threshold of <0.05 for selecting the significant pathways.

## Results

### CIGs Display Unique Genetic Sequence Signatures

Genetic sequence elements play important roles as cis-regulatory codes that dictate unique epigenetic patterns for the transcriptional regulation of CIGs. For example, super-enhancers tend to regulate CIGs and each comprises a long cluster of cis-regulatory elements that bind transcription factors(5, 68). The broad H3K4me3 pattern is also enriched at CIGs and tends to be associated with a broad distribution of transcription factor binding motifs(8). Therefore, we decided to perform a comprehensive survey of genetic sequence signatures together with expression features, expecting that some can be useful in uncovering CIGs.

Our meticulous analysis revealed 73 features enriched or depleted significantly in 247 CIGs, curated from literature(12), when compared to 245 control genes(12), 2142 housekeeping genes(69), or all human genes in the genome **(Figure 1a)**. Gene expression level is significantly higher for CIGs when compared to control genes and the entire human gene set, but intriguingly, shows little difference between CIGs and housekeeping genes **(Figure 1b)**. Gene expression specificity scores demonstrated a significant elevation in CIGs than in Housekeeping genes and other genes **(Figure 1c and S1a, b)**. This indicates that CIGs are overall more specific to certain cell types, consistent with their crucial role in maintaining unique cellular identities. Considering that expression specificity value for a gene will rely on the cell types used for comparison(70–72), and different CIGs display different degrees of expression specificity(12), we continued to explore genetic sequence signatures that might be useful to further improve accuracy for identifying CIGs. An investigation of the PhyloP100way score in the promoter regions of CIGs unveiled a higher degree of sequence conservation when compared to the other three gene categories, indicating stronger evolutionary conservation of CIG promoters across species **(Figure 1d and S1c, d)**. Moreover, CIGs exhibited stronger enrichment of binding motifs for transcription factors **(Figure 1e and Figure S1e)**, RNA-binding proteins **(Figure 1f and Figure S1e)**, and miRNAs **(Figure 1g and Figure S1e)**. CIGs displayed longer 3’-UTRs, potentially implicating greater involvement in post-transcriptional regulation, e.g., by miRNA binding sites, which are known as often occurring in 3’-UTRs (**Figure 1h**). This observation is consistent with the reported low RNA stability of CIGs(11), as miRNA medicates the decay of target RNAs in the cells(73, 74).

**Figure 1.**
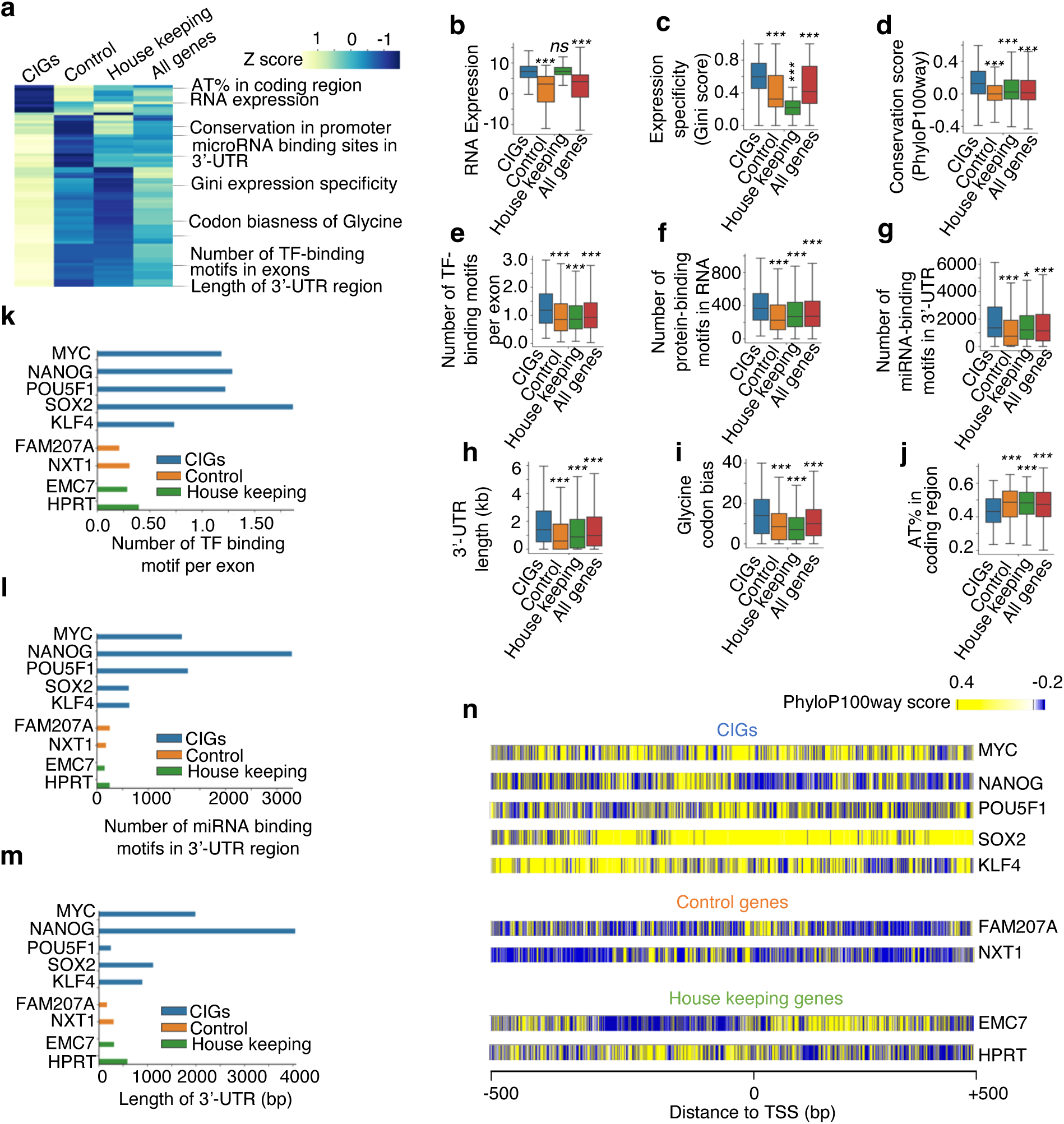
A systematic survey of genetic sequence signatures and RNA expression features for cell identity genes. **(a)** Heatmap illustrating the median values of 73 features that each displayed significant differences between cell identity genes and either control genes, housekeeping genes, or all human protein-coding genes. **(b-j)** Boxplots showing values of represent features in individual gene categories. P values determined by the two-tailed Wilcoxon test. *, p-value < 0.05; **, p-value < 0.01; ***, p-value < 0.001. **(k-m)** Bar plots showing genetic sequence feature values of individual embryonic stem cell identity genes and control or housekeeping genes. **(n)** Heatmap showing PhyloP100way scores around the Transcription Start Site (TSS) of individual embryonic stem cell identity genes and control or housekeeping genes.

Unexpectedly, the coding sequences of CIGs are longer than those of other gene categories, as shown in **Figure S1f**. We also found that there is a noticeable trend for the distance between the transcription start sites of CIGs and their nearest neighboring genes to be greater compared to other gene categories (**Figure S1g**). Additionally, a notable bias toward triplet codons encoding the amino acid Glycine was observed in CIGs (**Figure 1i**), suggesting a preference for loop-forming in protein structures(75) that facilitate protein interactions(76) and catalytic activities(77). Interestingly, CIGs exhibited a lower AT content in their coding regions **(Figure 1j and S1h)**, potentially associated with the known low RNA stability of CIGs(78, 79). Manual inspection confirmed these genetic sequence characteristics at many reported CIGs, e.g., the embryonic stem cell (ESC) identity genes MYC, NANOG, POU5F1, SOX2, and KLF4 but not at the housekeeping genes HPRT and EMC7 **(Figure 1k-n)**. It will be interesting to investigate the biological mechanisms underlying the association between these genetic signatures and CIGs.

### SCIG Uncovers CIGs Accurately by Genetic Signatures and Expression Information

Motivated by the observed significant difference in genetic signatures and expression patterns between the known CIGs versus the control gene categories, we developed SCIG, a logistic regression-based machine learning model to uncover new CIGs.

The significant features were obtained through the Wilcoxon test, followed by removing the multicollinearity among the features. This procedure yielded 73 features out of a comprehensive list of 680 candidate features **(Figure S2a)**. We next further performed forward feature selection to determine the optimal subset of these features for the logistic regression model **(Figure S2a)**. Evaluation metrics such as Matthew’s correlation coefficient and F1-score indicated the superior predictive performance of features selected by this pipeline when compared to conventional feature selection methods, including the Select K, ANOVA F1-score, mutual information, forward, backward, and recursive feature elimination (RFE) **(Figure S2b and S2c)**. By increasing the feature number starting from 1, the model with more features shows better performance but requires up to 19 features to achieve superior performance **(Figure 2a)**, as a further increase in feature number does not significantly improve the performance **(Figure S2d)**. The 19 key features include RNA expression level, expression specificity scores (Gini and Tau matrices), codon biases, sequence conservation, number of transcription factor binding motifs, number of protein-binding motifs in RNA, number of miRNA binding motifs, nearby promoter distance, etc. (**Figure 2b**). These features accurately distinguished between known CIGs and control genes with an area under the receiver operating characteristic curve (AUROC) of 0.95. Integration of these 19 features, SCIG successfully recaptured the known CIGs with an accuracy of 88.8%, sensitivity of 91.4%, and specificity of 89.3% **(Figure S2e)**. RNA expression-based features, specifically the RNA expression-level and expression-specificity matrices, exhibited stronger feature coefficients when compared to the genetic sequence features **(Figure 2b)**. This observation is consistent with results from the leave-one-out feature test strategy **(Figure S2f)**. However, integrating the genetic sequence signatures resulted in a model significantly better than models using only the expression information (**Figure 2a**).

**Figure 2.**
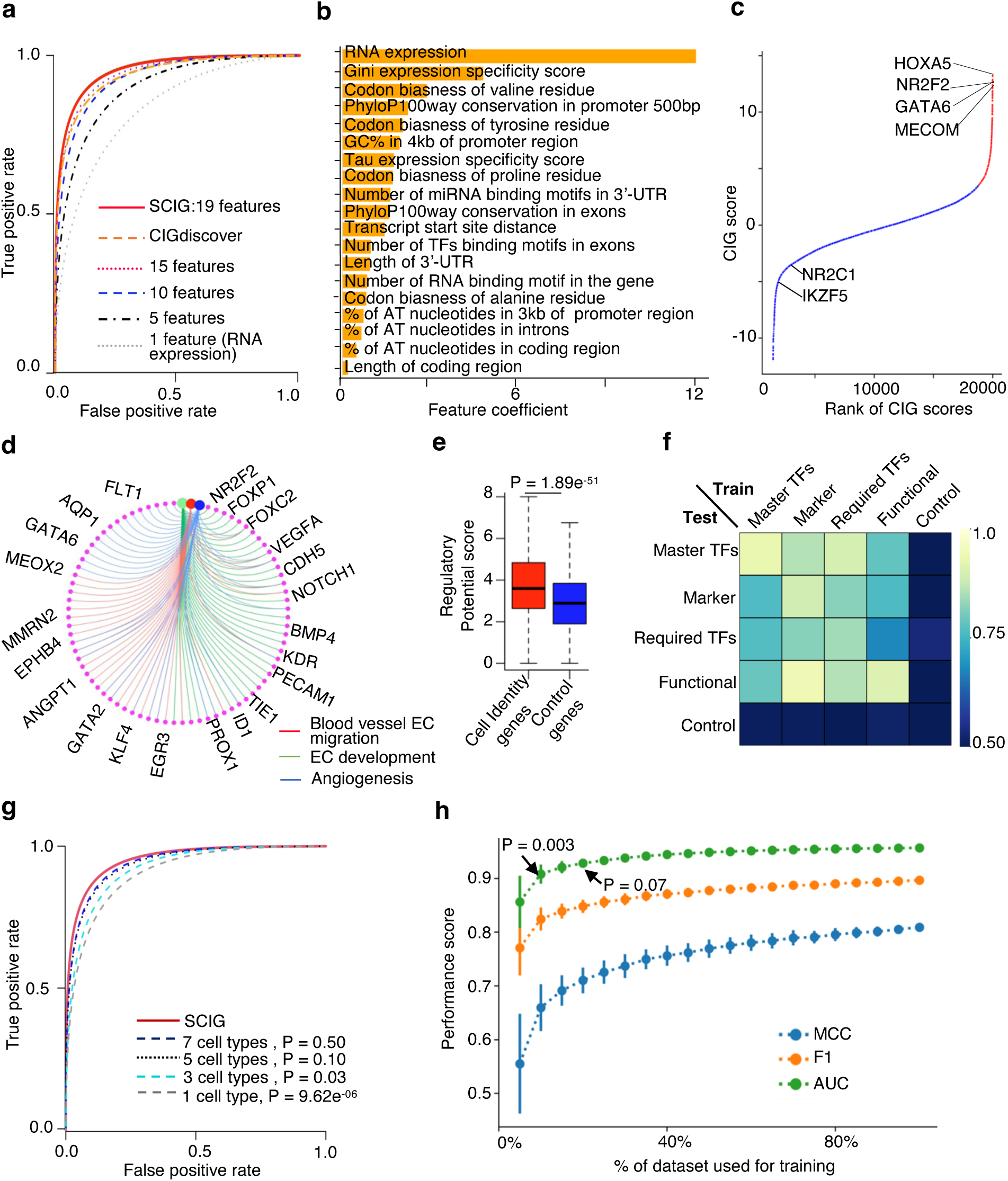
SCIG combines genetic sequence signatures with expression information to uncover cell identity genes in a cell. **(a)** ROC curve illustrating the performance of SCIG with varying numbers of top features. Performance of the CIGdiscover algorithm that uncovers CIGs by histone modification signatures and gene expression information is also presented. **(b)** Bar plot showing feature coefficient of individual genetic sequence signatures or gene expression features used for machine learning in SCIG. **(c)** Rank plot presenting the CIG scores of individual genes in HUVEC. Red and blue colors indicate CIGs and other genes defined by SCIG respectively. **(d)** The CIGs defined by SCIG in HUVEC are enriched with endothelial pathways. **(e)** Boxplot of regulatory potential scores demonstrating the regulation intensity of individual genes by transcription factors in HUVEC. **(f)** Heatmap showing AUROC of SCIG variants trained and tested by individual gene categories. **(g)** ROC curves depicting the performance of SCIG, and its variants trained with data from varying numbers of cell types. **(h)** Line plot illustrating the performance of SCIG, and its variants trained with varying subsets of the known CIGs.

The default algorithm in SCIG is a logistics regression model and achieved remarkable performance, with a Matthews correlation coefficient of 0.80 (**Figure S2b**) and an F1-score of 0.90 (**Figure S2c**) in recapturing the known CIGs. For comparison, we assessed the predictive capacity of alternative machine-learning algorithms. The logistic regression model exhibited a lower error rate and a higher Matthews correlation coefficient, underscoring its superior performance compared to the other models, including the Naive Bayes, support vector machines, and AdaBoost (**Figure S2g**). Applying SCIG to human umbilical vein endothelial cells (HUVECs), well-known endothelial cell identity genes such as the NR2F2(80, 81), MECOM(82), HOXA5(83), and GATA6(84) were top-ranked by SCIG, indicating the ability to recapture the endothelial CIGs **(Figure 2c)**. The identified CIGs are enriched with endothelial pathways, reaffirming their association with EC identity **(Figure 2d and Figure S2h)**. As CIGs tend to be regulated by super-enhancers(5, 68), we interrogated ChIP-seq profiles of transcription factors from the Cistrome browser(85). The result verified that the predicted CIGs exhibited a pronounced enrichment of transcription factor binding events compared to control genes **(Figure 2e)**. Additionally, we extended the SCIG model to the mouse genome, achieving an AUROC of 0.94 **(Figure S3a)** utilizing 16 distinct genetic and RNA expression features **(Figure S3b)**. SCIG consistently identified pan-endothelial cell marker genes as CIGs in brain, lung, and liver endothelial cells **(Figure S3c-e)**. These results indicate that the algorithm in SCIG is optimal for accurately recapturing known CIGs based on the combination of genetic sequence signatures and RNA expression patterns.

### Robust Performance of SCIG with Small and Noisy Training Data

We assessed whether the performance of SCIG is consistent when applied to four CIG categories defined based on literature curation in recent work(12). These include master transcription factors whose expression is reported as sufficient to induce a cell type, required transcription factors whose depletion is reported to impair the differentiation toward a cell type, function genes reported to play a cell type-specific function, and marker genes that are simply used as marker of a cell type in literature. The model consistently demonstrated robust performance (AUROC > 0.83) when applied to each of these categories **(Figure S4a)**. Impressively, a model trained by one CIG category can accurately recapture the other CIG categories but not the control genes, underscoring the similarity in characteristics of these CIG categories **(Figure 2f)**.

To test whether the size of the training data has been large enough to reach an optimal performance, we evaluated the algorithm trained with different subsets of the training data. The algorithm’s performance improved when the known CIGs and control genes from more cell types were used to train the model, with the improvement saturated at up to 5 cell types (**Figure 2g**). Also, the model requires up to 15% of the 247 known CIGs to achieve optimal performance (**Figure 2h**). Considering the potential imbalance between the number of control genes and CIGs in a cell, we investigated the impact on SCIG performance. We systematically varied the number of non-identity genes used to train the model and analyzed its effect on SCIG accuracy. Despite increasing the number of non-identity genes up to 20-fold higher than the number of CIGs, we observed minimal changes in SCIG performance **(Figure S4b)**.

We investigated whether the model trained by known CIGs from one cell type can accurately recapture the known CIGs of a different cell type. To this end, we performed two different tests for comparison. We first performed a parallel test, where the model was trained and tested based on known CIGs from the same cell type. We then performed a cross-test, where the model was trained by genes from one set of cell types and tested by genes from another set of cell types. The results revealed that the algorithm performance is excellent in both the parallel test (AUROC 0.93) and the cross-test (AUROC 0.90) **(Figure S4c)**.

We extensively evaluated the robustness of SCIG against noise in the training data. To this end, the labels of a randomly picked subset of the known CIGs and control genes were swapped. As expected, the algorithm performance decreased along with the increased noise ratio in the training data. However, with 20% mislabeled CIGs or control genes, the AUROC only decreased from 0.95 to 0.75 **(Figure S4d)**, suggesting that the algorithm is considerably resilient to noise in the training data.

### SCIGNet Uncovers Master Transcription Factors of Cell Identity Genes by Machine-learning Analysis of Network Features

It is well recognized that a cell identity can be established by a small cocktail of master transcription factors, such as the Oct4, Sox2, Klf4 and c-Myc for pluripotent stem cells(4, 5). As a cell type often has several hundred or more CIGs, we next developed another logistic regression model, SCIGNet, to uncover the master transcription factors of a cell identity based on the network features of CIGs. Conventional network analysis often defines master transcription factors based on their number of downstream target genes. However, it is unclear if there are other network features that might be more useful to define the master transcription factors of a cell identity. To this end, we considered a set of 23 network features, including the number of downstream as well as upstream network edges connected to a transcription factor, gene expression values, CIG scores, etc. (**Figure 3a).** SCIGNet learns the features associated with the known master transcription factors of cell identity curated from literature(12). It then optimizes the weight per feature to combine these features for predicting new master transcription factors.

**Figure 3.**
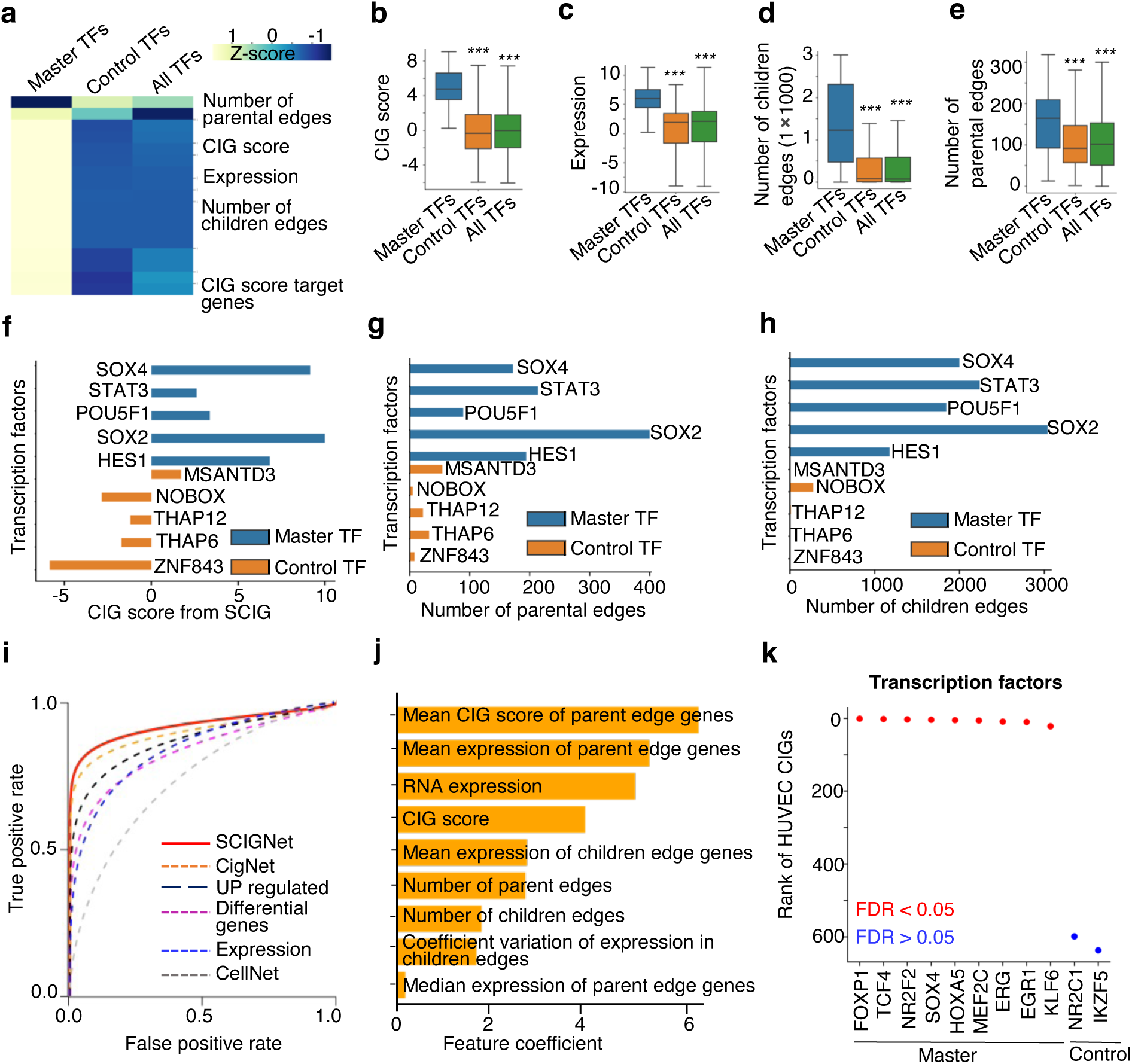
SCIGNet combines network features of cell identity genes to uncover master transcription factors of a cell identity. **(a)** Heatmap showing Z scores of individual network feature values for individual transcription factor groups. TF, transcription factor. **(b-e)** Box plot illustrating representative network feature values for individual transcription factor groups. *, p-value < 0.05; **, p-value < 0.01; ***, p-value < 0.001. **(f-h)** Bar plot showing feature values of individual transcription factors. **(i)** ROC curves showing the performance of SCIGNet and other methods for uncovering master transcription factors of cell identity. **(j)** Bar plot showing feature coefficients of individual network features used by the machine learning models in SCIGNet for uncovering master transcription factors. **(k)** Rank of individual transcription factors based on the scores calculated by SCIGNet in HUVEC.

Our feature analysis revealed that reported master transcription factors exhibit greater CIG scores calculated by the SCIG **(Figure 3b)** and higher expression values **(Figure 3c)** compared to other transcription factors. Additionally, when compared to other transcription factors in the CIG network, we observed a greater number of children edges connecting master transcription factors to downstream target genes **(Figure 3d)**. This observation is consistent with the role of these factors as a master regulator in the network. Intriguingly, we also observed a greater number of parental edges connecting the master transcription factors to their upstream regulators (**Figure 3e**). This observation indicates that the master transcription factors themselves, when compared to other transcription factors in the CIG network, are under more regulations. These results were also observed by manually inspecting the feature values of individual master transcription factors in the embryonic CIG network (**Figure 3f-h**).

The SCIGNet model employed 9 features to achieve optimal performance with an AUROC of 0.91 **(Figure 3i)**. Unexpectedly, the mean CIG score and average RNA expression of the genes connected by the parental edges appeared to be the two most useful features for identifying the master transcription factors, followed by the RNA expression and CIG score of the master transcription factors themselves **(Figure 3j)**. When compared to simple methods based on analysis of gene expression level or the comprehensive network model CellNet, SCIGNet always showed a better performance in recapturing the known master transcription factors **(Figure 3i)**. Furthermore, we employed SCIGNet in HUVEC cells to pinpoint master transcription factors regulating their identity. We uncovered that most of the top-ranked transcription factors, such as FOXP1(86), NR2F2(80), SOX4(87), MEF2C(88), and ERG(89), have already been experimentally characterized for their role in the regulation of HUVEC identity **(Figure 3k)**. Also, applying the model to mice cells identifies the master transcription factors in the CIG network with an AUROC of 0.93 (**Figure S3f**).

### CIG Score Outperforms Expression Value in Capturing Cell Identity in Network Analysis

We next tested if the CIG scores calculated by SCIG can be better than the gene expression values when aiming at uncovering CIG networks in single-cell RNA-seq analysis. We used SCIG to calculate a CIG score for each gene in each single cell based on single-cell transcriptomes of healthy human fetal hearts at 22 weeks(58) **(Figure 4a)**. We next clustered cells using either the CIG scores or the RNA expression values based on highly variable genes. Both methods effectively recaptured the 12 cell types, including cardiomyocytes, endothelial cells, fibroblast, mesothelial cells, proliferating cells, mast cells, erythroblasts, etc., as reported in the original study (**Figure 4b)**. Thereafter, cell clusters from the two methods were independently used as input for network analysis based on either the expression values or CIG scores using the hdWGCNA package(59). Employing gene expression data, we obtained six network modules (referred to as the expression network), while the CIG score yielded nine modules (referred to as the cell identity score network). The specificity of hub genes towards particular cell types was illustrated in a dot plot, showcasing their expression **(Figure 4c left)** or CIG score **(Figure 4c right)**.

**Figure 4.**
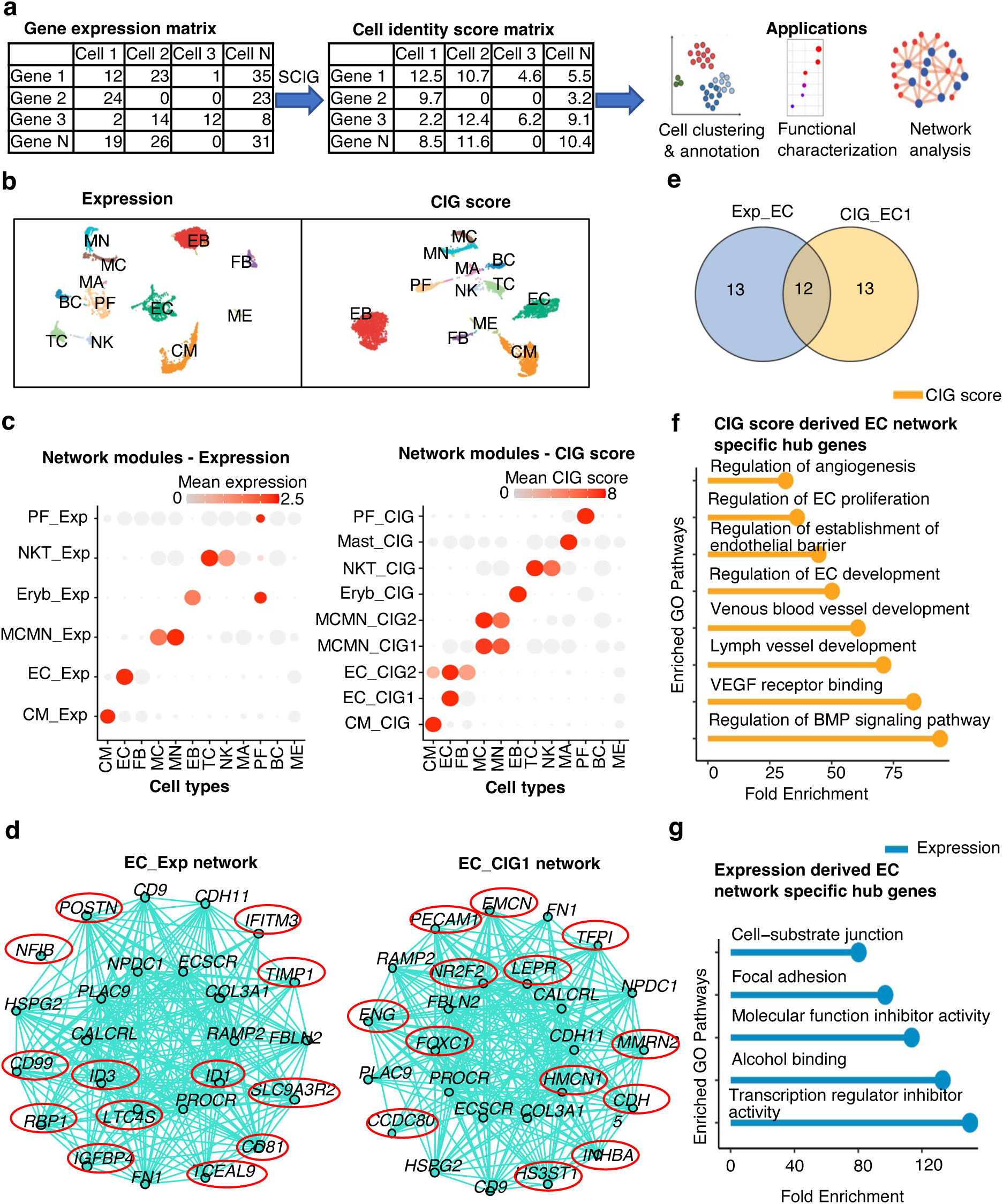
CIG score outperforms expression value in capturing cell identity in network analysis. **(a)** Workflow to use SCIG for uncovering cell identity genes at the single-cell level and perform subsequent applications. **(b)** Single-cell clustering based on expression and CIG score. **(c)** Network modules defined using hdWGCNA based on expression and CIG score matrices. **(d)** Network plot showing the top 25 hub genes in endothelial cell identity score- and gene expression-derived network. Expression- and CIG score-specific network hub genes are marked in red circles. **(e)** Ven diagram showing overlap of top 25 hub genes in CIG score- and gene expression-derived networks. **(f)** Pathway enrichment analysis of CIG score-specific network hub genes. **(g)** Pathway enrichment analysis of expression-specific network hub genes. CM, cardiomyocytes; EC, endothelial cells; FB, fibroblast; MC, macrophages; MN, monocytes; EB, erythroblast; TC, T cells; NK, natural killer cells; MA, mast cells; PF, proliferating cells; BC, B cells; ME, mesothelial cells.

For a fair comparison between the expression network and cell identity score network for a cell type, we focused on the top 25 hub genes from each network. For endothelial cells (EC), a cell identity score network (EC_CIG1) and the expression network (EC_Exp) **(Figure 4d)** shared 12 hub genes **(Figure 4e)**. Notably, pathway analysis revealed that EC cell identity score network-specific hub genes were associated with the regulation of EC and blood vessel development, VEGF signaling, and angiogenesis **(Figure 4f)**. In contrast, expression network-specific hub genes were associated with housekeeping-related cellular functions **(Figure 4g)**. This observation suggests that the CIG score is superior in revealing the EC identity network. Interestingly, based on CIG scores, hdWGCNA further constructed a network module shared by endothelial cells, fibroblasts, and cardiomyocytes (EC_CIG2) **(Figure 4c right and S5a)**. Comparing this network with the EC_Exp module revealed only 2 common hub genes (**Figure S5b**). Pathway analysis indicates that the genes in the EC_CIG2 model are involved in the well-known epithelial (endothelial) to mesenchymal transition (**Figure S5c**), which plays critical roles in the development and many diseases, such as heart failure(90, 91).

For the cardiomyocyte network modules, the expression network **(Figure S5d)** and cell identity score network **(Figure S5e)** shared a set of 10 hub genes **(Figure S5f)**. The hub genes specific to the cell identity score network demonstrated a more pronounced enrichment across pathways regulating the cellular processes and functions specific to cardiomyocytes **(Figure S5g)**. The network modules derived from expression **(Figure S6a)** and cell identity score **(Figure S6b)** also uncovered different hub genes for the proliferating cell population. Upon pathway enrichment analysis, it became evident that the cell identity score network hub genes displayed a heightened association with cell cycle-related pathways (**Figure S6d**), contrasting with expression networks, which are only enriched with some housekeeping pathways (**Figure S6c**). Two network modules, MCMN_CIG1 and MCMN_CIG2, were identified as common for macrophages and monocytes using cell identity scores, while one module, MCMN_Exp, was identified based on expression values **(Figure 4c)**. We observed that 18 hub genes were shared between the MCMN_CIG2 and MCMN_Exp modules **(Figure S6e)**. However, there were no overlapping hub genes between the MCMN_CIG1 and MCMN_Exp modules. Notably, the genes in the MCMN_CIG1 cell identity network (**Figure S6f**) were enriched with pathways related to inflammatory response and cellular response **(Figure S6g)**. With the expression values, there was no network module detected for mast cells. Conversely, the unique cell identity score network of the mast cells unveiled an enrichment pattern aligning with mast cell functions, including regulation of chemokine ligand production and immune responses (**Figure S6h**)(92, 93). Meanwhile, for erythroblast and natural killer T cells, less difference was observed between the expression- and CIG score-derived networks (**Figure S6i-j**), with no major distinction concerning their pathway associations. Overall, we posit that networks derived from CIG scores prove more advantageous in elucidating the gene network involving cell identity regulation.

### Cell Identity Score Improved Single-cell Trajectory Analysis of Neuronal Differentiation

We harnessed the SCIG method to explore the landscape of CIGs in 1720 single-cells during human forebrain glutamatergic neuron differentiation. This dataset effectively captured the dynamic process of mature neuron formation starting from radial glial progenitors, exhibiting a linear trajectory(60, 94). We performed single-cell clustering using either the RNA expression values or the CIG scores based on top 2000 high-variation genes. There was only a 24% overlap between the highly variable genes defined by these two methods (**Figure 5a**). The highly variable genes identified through the CIG score exhibit a stronger enrichment in the forebrain and neuronal development pathways (**Figure 5b).** Meanwhile, the clustering based on CIGs recapitulated all the cell types defined in previous studies based on gene expression(60, 94)(**Figure 5c-d, Figure S7a-b)**. However, the Silhouette coefficient, an internal cluster validation measure, is 3 folds smaller for the CIG-based than for the expression-based methods (**Figure S7c**). This suggests that the CIG scores are better at capturing the relation between the cells in the differentiation process, as also can be observed in the single-cell clustering architecture derived from highly variable genes identified uniquely by expression or CIG score (**Figure S7b**). Meanwhile, we found substantial differences in the sizes of cell populations defined based on CIG score- versus gene expression-based clustering, e.g., population sizes of Neuroblast1, Immature neurons, and mature neurons (**Figure 5e**). The greatest cell identity switch between the two methods happened to neuroblast1 and immature neuronal cells (**Figure 5f**).

**Figure 5.**
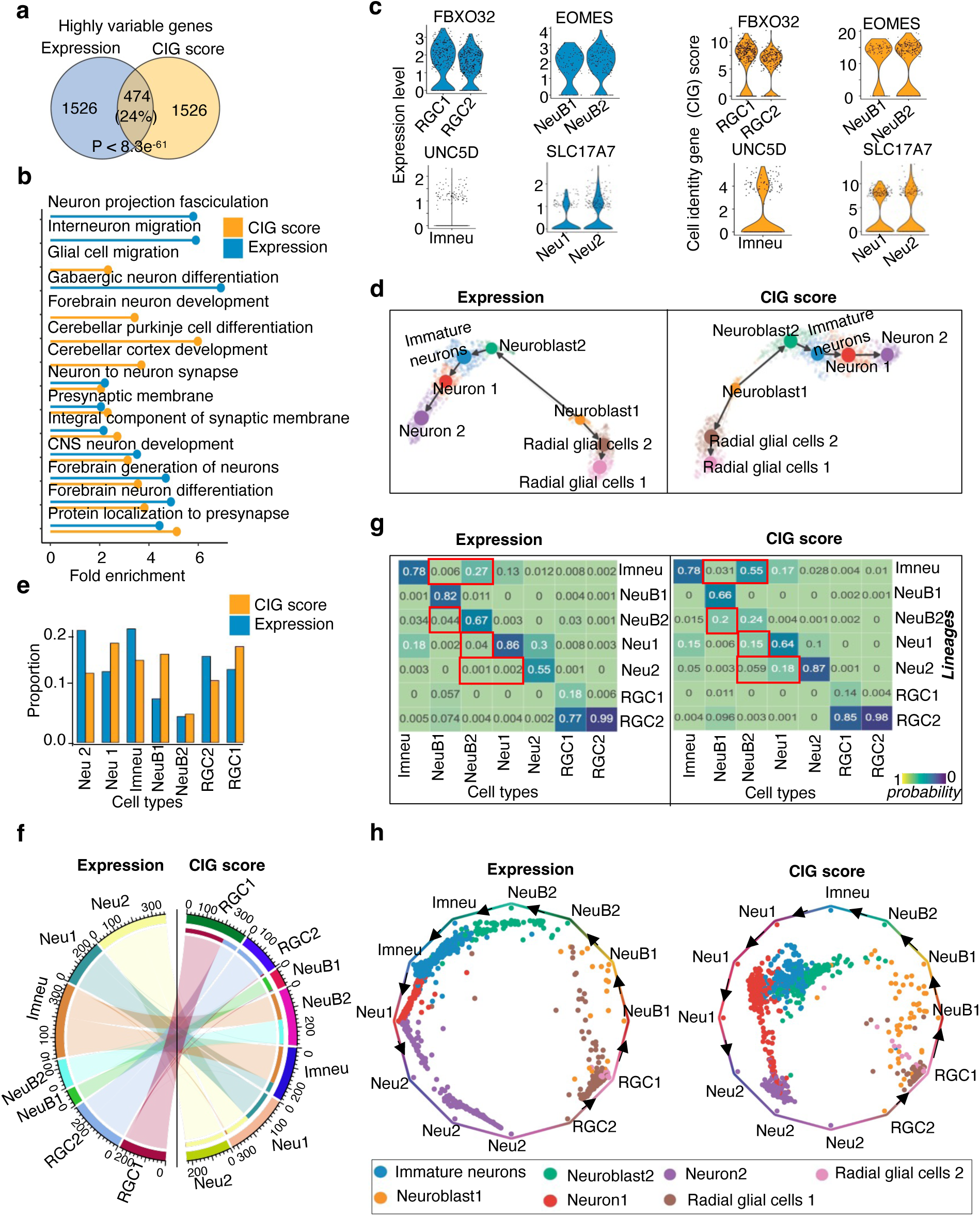
Cell identity score improved single-cell trajectory analysis of neuronal differentiation. **(a)** Venn diagram showing overlap between highly variable genes defined based on single-cell gene expression and CIG scores. **(b)** Pathway enrichment analysis of highly variable genes defined based on single-cell gene expression and CIG scores. **(c)** Expression level and cell identity score of marker genes that we used to define the cell types in this dataset. **(d)** UMAP displaying cell types and differentiation trajectories in the human forebrain glutamatergic neurogenesis dataset. **(e)** Barplot showing proportions of individual cell populations clustered based on CIG scores or expression values. **(f)** Chord diagram showing the cells that are switched between expression- and CIG score-based cell clustering. **(g)** Heatmaps depicting transition probabilities quantified using CellRank between cell populations clustered based on expression values (left) or CIG scores (right). **(h)** Projection plots showing the fate probabilities of each cell during the glutamatergic neuron genesis trajectory. RGC1, radial glial cells 1; RGC2, radial glial cells 2; NeuB1, Neuroblast1; NeuB2, Neuroblast2; imneu, immature neurons; Neu1, Neuron 1; Neu2, Neuron 2.

To assess the effects of cell type rearrangements between the two methods on differentiation trajectory analysis, we conducted RNA velocity analysis and quantified the transition probabilities between the cell populations using the algorithm CellRank(63). Although the result indicated an overall consistent neuronal differentiation trajectory between the two methods (**Figure 5d**), the probability of the expected transition from neuroblast 1 and neuroblast 2 towards immature neurons was increased by two to five-fold by the CIG-based clustering compared to the expression-based one (**Figures 5g**). Similarly, the transition probability from neuroblast 1 to neuroblast 2, neuroblast 2 to immature neuron, and neuron 1 to neuron 2 also increased in the CIG-based clustering result (**Figure 5g**). In contrast, the probability of transition that is unknown or opposite to literature report, e.g., from neuroblast 2 to neuroblast 1 and neuron 2 to neuron 1, exhibited decreases (**Figure 5g**). We next presented the transition probabilities of each cell using a circular projection plot, positioning naive or intermediate cells at the center, while mature or fate-biased cells fell in the corners corresponding to their respective identities. This visualization highlighted the relationships between cell types during differentiation. From the cell types derived using CIG scores, we observed that intermediate cell types, such as immature neurons, tend to cluster closer to the middle of the plot, which is better than the gene expression method (**Figure 5h)**. In contrast, matured or terminated differentiated cell types, including radial glial cells and neurons, are situated at the corners corresponding to their respective identities. These results suggest that single-cell clustering based on CIG scores can better arrange cells in the differentiation trajectory.

### SCIG Recapitulated the Landscape of CIGs in the Endothelial Differentiation Process

We further applied the SCIG algorithm to identify CIGs in the scRNA-seq profiles from various stages during the differentiation of human embryonic stem cells (H9) into endothelial cells(64), including stem cells (day0), mesoderm (day4), EC-mesenchymal progenitors (day 6 and 8), and endothelial cells (day12) (**Figure S8a)**. Subsequent pathway enrichment analysis of the identified CIGs at each stage revealed stage-related pathways (**Figure S8b-e)**. To investigate the relationship between expressed genes and identified CIGs, we compared the top 10% highly expressed genes with the top 10% cell identity genes and observed only a 26% overlap (**Figure 6a)**. Pathway enrichment analysis showed that the unique high-scored CIGs are enriched with EC functional pathways, including VEGF signaling, BMP signaling, and endothelium development. In contrast, the highly expressed genes are involved in housekeeping-related functions such as RNA splicing and ribonucleoprotein complex processes (**Figure 6b)**. Pearson correlation analysis between the CIG scores and the expression levels of highly expressed genes revealed a minimal correlation, which is reasonable because the CIG score combines expression information and genetic sequence signatures (**Figure S8f and S8g)**. Therefore, the CIG score is better than the expression level for enriching cell identity pathways.

**Figure 6.**
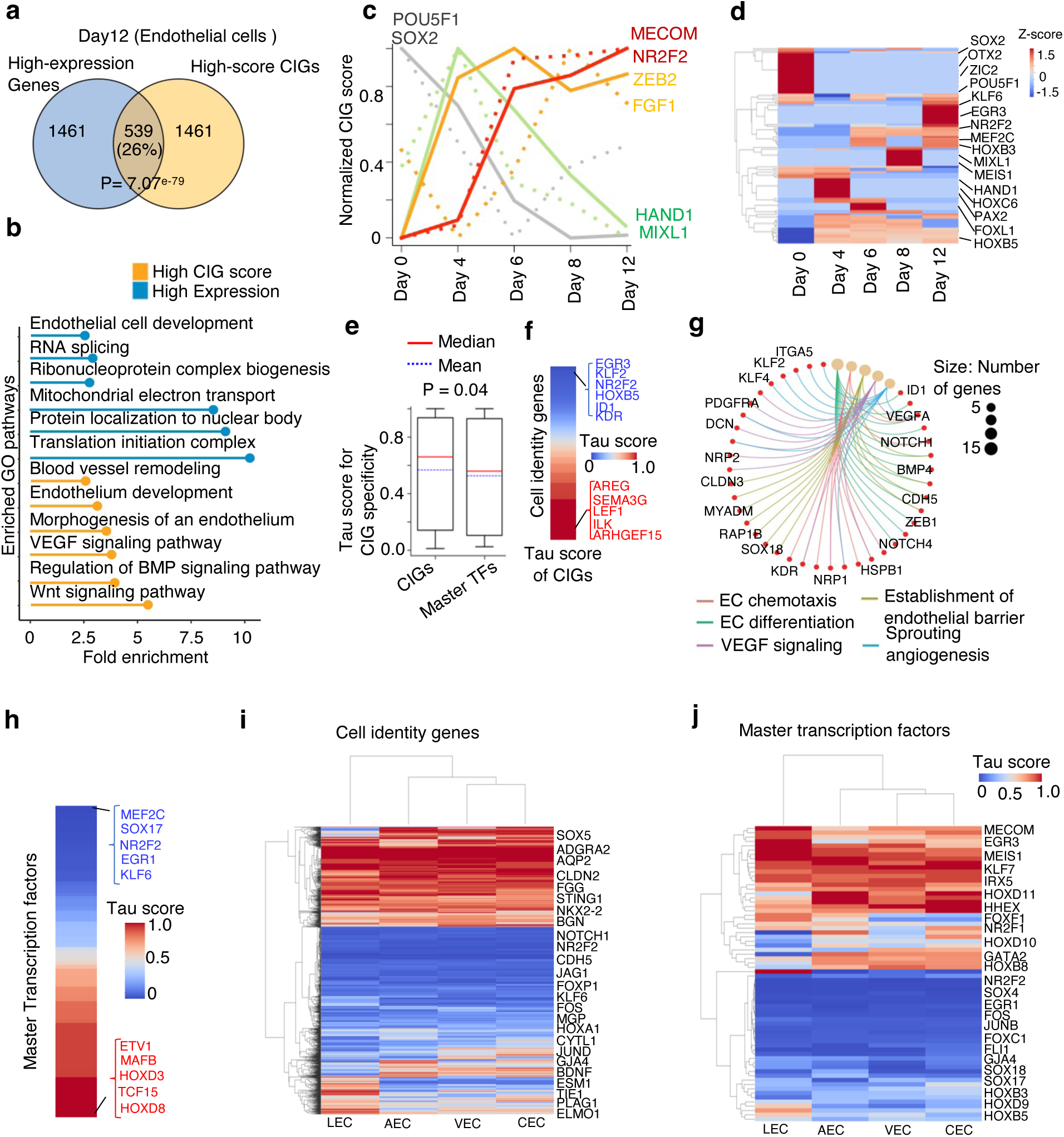
SCIG revealed new insight into ec identity fine-tuning by tissue microenvironment. **(a)** Venn diagram illustrating overlap between the top 10% highly expressed genes and top 10% high-score cell identity genes identified by SCIG in endothelial cells. **(b)** Pathway enrichment analysis of the top 10% highly expressed-specific genes and top 10% high-score cell identity specific genes identified by SCIG in endothelial cells. **(c)** CIG scores for known marker genes of ESC (SOX2, POU5F1), Mesoderm (MIXL1, HAND1), EC-mesenchymal progenitors (FGF1, ZEB2), and endothelial (MECOM, NR2F2) cells during the ESC to EC differentiation process. **(d)** Heatmap showcasing the identified master transcription factors of CIGs across different stages of ESC to EC differentiation. **(e)** Box plot showing Tau score of CIGs and their master transcription factors uncovered for endothelial cells across 15 tissue types. **(f)** Heatmap showing the Tau score of endothelial CIGs. **(g)** Gene-concept network plot displaying pathways enriched in the endothelial CIGs conserved across 15 tissue types. **(h)** Heatmap showing the Tau score of endothelial master transcription factors of CIGs. **(i)** Heatmap showing Tau score of CIGs in each of the four EC subtypes across 15 tissue types. **(j)** Heatmap showing Tau score of CIG master transcription factors in each of the four EC subtypes. Data for arterial EC (AEC), venous EC (VEC), capillary EC (CEC) and lymphatic EC (LEC) were presented.

SCIG exhibited the ability to identify CIGs that are specific to individual stages. These include the SOX2(4), and POU5F1(OCT-4)(4) for stem cells (Day 0), the HAND1(95), and MIXL1(96) for mesoderm cells (Day 4), FGF1(97) and ZEB2(98) for EC-mesenchymal progenitor cells (Day 6 and 8), and the MECOM(82), and NR2F2(80) for EC (Day 12) (**Figure 6c)**. Meanwhile, notable overlap of CIGs between neighboring stages were also recapitulated, e.g., between stem cells (Day 0) versus mesoderm cells (Day4) stages and EC-mesenchymal progenitors (Day 6, 8) versus ECs (Day 12) stages (**Figure S8a)**. Additionally, we used the SCIGNet to identify the network regulators of CIGs. Consistent with the literature report, we observed that the SOX2(4), POU5F1(4), and OTX2(99), served as key regulators in stem cells, PAX2(100), HOXC6(101) and HAND1(95) in mesoderm cells, while the NR2F2(81), KLF6(102) and EGR3(103) are key regulators in endothelial cells (**Figure 6d)**. Therefore, SCIGNet successfully recapitulated the network regulators governing the differentiation process of human embryonic stem cells toward the endothelial cell fate.

### SCIG Revealed New Insight into EC Identity Refinement by Tissue Microenvironment

We obtained single-cell transcriptomes representing ECs from 15 tissue types in the DISCO database(65), including the adipose, bladder, breast, gut, heart, intestine, kidney, liver, lung, ovary, skeletal muscle, skin, stomach, testis, and thymus tissues. The EC in each tissue type comprises 4 EC subtypes, including the arterial, capillary, venous, and lymphatic ECs. We used SCIG to elucidate the identity gene landscape in each EC subtype from each tissue type. The algorithm identified a total of 2067 CIGs, including 86 CIG master regulators. For each gene, we computed a Tau score representing cell identity score specificity across all tissues, ranging from 0 to 1. A Tau score close to 0 and 1 indicates low and high tissue specificity, respectively.

Intriguingly, we found CIGs tend to exhibit significantly greater Tau scores than CIG master transcription factors **(Figure 6e).** This difference between CIGs and their master transcription factors is consistent when we further analyze each EC subtype across the tissue types, with the greatest difference observed for venous EC followed by lymphatic EC (**Figure S9a-d**). The Genes such as EGR3(103, 104), KLF2(105, 106), NR2F2(80, 81), HOXB5(107), ID1(108), and KDR(109) possess moderate Tau scores, indicating conservation across tissue types, while AREG(110), SEMA3G(111), LEF1(112), ILK(113), and ARHGEF15(114) exhibit greater Tau scores **(Figure 6f)**. The conserved CIG genes are enriched with EC-related biological processes and functions (**Figure 6g**). In contrast, the low-conservation CIGs demonstrate weaker enrichment in EC functions and a greater association with tissue-specific functions (**Figure S9e**). Expanding this analysis to EC master transcription factors, MEF2C(88), SOX17(115), NR2F2(116), EGR1(117), and KLF6(102), appeared conserved across tissue types, while MAFB(118), ETV1(119), TCF15(120), and HOXD8(121) appeared to be less conserved (**Figure 6h**). These results imply that the tissue microenvironment, as an additional factor to the master transcription factors in the cells, may play an important role in fine-tuning the EC identity.

Meanwhile, there are also a few differences in CIGs between endothelial subtypes **(Figure 6i)**. The conserved CIGs of each EC subtype across the tissue types include the GJA4(122), CXCL12(123, 124), and NOTCH4(125) in arterial EC (**Figure S9f)**, the NR2F2(80), NRP2(126), and EPHB4(127) in venous EC (**Figure S9g)**, the FABP5(128), SPARC(129), and CD36(130) in capillary EC (**Figure S9h)**, and the PROX1(131), LYVE1(132), and CCL21(133, 134) in lymphatic EC (**Figure S9i**). On the other hand, we also observed tissue-specific CIGs for each EC subtype. For example, the VEGFC(135), MYCN(136), and EFNB2(137) in arterial EC (**Figure S9f)**, the FZD5(138), ARG1(139), and LRG1(140) in venous EC (**Figure S9g)**, the VTN(141), RGCC(129), and PRX(142) in capillary EC (**Figure S9h)**, and the CCL4(143), FOXO3(144), and MMRN2(145) in lymphatic EC (**Figure S9i)**. In the case of master transcription factors **(Figure 6j)**, the algorithm identified SOX17(146) and KLF6(102, 147) in arterial EC (**Figure S9j)**, NR2F2(80, 148) in venous EC (**Figure S9k)**, MEF2C(149) in capillary EC (**Figure S9l)**, and NR2F1(150) and NR2F2(151) in lymphatic EC (**Figure S9m**) as conserved across the tissue types. These analyses shed light on the heterogeneity of endothelial CIGs across tissue types, aiding our understanding of tissue-specific microenvironments in cell identity refinement.

## Discussion

Advancements in high-throughput sequencing technologies, particularly scRNA-seq, have revolutionized the study of gene expressions at a large scale. This technique has enabled researchers to delve into cellular heterogeneity, cell fate conversion, and other cellular processes. Investigating the role of genes in cellular identity regulation at the single-cell level is crucial for gaining insights into cell fate conversion and its potential applications in regenerative medicine.

The community has been analyzing CIGs by considering differences in epigenetic modification patterns between CIGs and other genes in the same cell type, or by considering differential expression between cell types. If one could define CIGs solely based on gene expression in a cell type without comparing to other cell types, it would offer great advantages. For example, it will be straightforward to implement, making it suitable for large-scale analyses using bulk as well as single-cell expression profiles. However, relying solely on gene expression-based analysis may not yield optimal results, as distinguishing between CIGs and other expressed genes, including housekeeping genes and genes expressed in response to some conditions, can be challenging. Epigenetic modification profiles are useful because CIGs tend to display unique modification patterns that do not enrich other gene categories. However, profiling epigenetic modifications at a single-cell level represents a current technological challenge. The abundance of available single-cell expression data greatly surpasses that of epigenetic data, making gene expression-based approaches more accessible. Thus, we proposed a solution that combines gene expression information with genetic sequence signatures, which overcomes the drawbacks of strategies that rely on expression data alone or in combination with epigenetic profiles.

The utilization of cis-regulatory codes in genetic sequence has proven valuable in predicting transcriptional regulation(27, 152), 3D genome organization(153), RNA structure(24), and more. In our study, we employed a comprehensive set of genetic sequence signatures, including the PhyloP100way conservation score, transcription factor binding motifs, protein-binding motifs in RNA, miRNA-binding motifs, codon biases, and gene architectural features, to characterize CIGs. Our analysis revealed intriguing genetic sequence patterns in CIGs compared to other genes. Specifically, CIGs exhibited greater sequence conservation in their promoters, probably implying the importance of the conserved sequence in a tight regulation of transcription. Additionally, we observed that CIGs are enriched with the binding sites of a large number of transcription factors and miRNAs, consistent with the reported frequent transcription elongation(8) and low RNA stability of CIGs(11). Furthermore, the 3’ UTR regions of CIGs appeared longer than those of other genes, indicating a propensity to recruit regulatory factors, such as miRNA. These findings underscore the significance of cis-regulatory codes in DNA sequence for characterizing CIGs and shed light on the potential genetic codes that govern cell identity. Leveraging this information, we developed the SCIG algorithm, which identifies CIGs by combining RNA expression information with these genetic signatures. This novel strategy enables the prediction of CIGs at both the single-cell and bulk levels based on easily accessible RNA-seq and genetic sequence data. Applying SCIG to HUVEC cells successfully uncovered experimentally verified endothelial CIGs. Utilizing the CIGs identified by SCIG and gene regulatory network information, we further developed the SCIGNet algorithm that analyzes the expression regulation networks of CIGs to define master transcription factors governing a cell identity.

By applying SCIG to human fetal heart single-cell data and using the CIG score for cell clustering, we effectively recapitulated cell types identified by gene expression analysis. Moreover, network analysis for each cell type using hdWGCNA revealed that CIG score-derived network hub genes align with cell identities. In contrast, expression-derived network hub genes, e.g., for endothelial cells and proliferating cells, are more likely to enrich housekeeping-related functions. This result suggests the superiority of the CIG score for constructing cell identity gene network. On the other hand, in the human neurogenesis dataset, the CIG score more effectively organizes cells during single-cell clustering, which reflects differentiation trajectory better when compared to expression-based analysis.

We further applied SCIG to explore the spectrum of CIGs during the differentiation from human embryonic stem cells to endothelial cells, recapitulating stage-specific CIGs. For instance, SOX2(4), and POU5F1(OCT-4)(4) for embryonic stem cells, HAND1(95) and MIXL1(96) for mesoderm cells, and NR2F2(80) and MECOM(82) for endothelial cells were recapitulated. Similarly, exploring the endothelial CIG landscape across 15 tissue types revealed putative CIGs that are either conserved across tissue types or tissue-type specific. Conserved CIGs exhibited stronger enrichment in endothelial cell-related functions, while specific CIGs were weakly associated with EC-related pathways but significantly enriched in tissue-specific functions. Intriguingly, the tissue specificity appeared to be greater for CIGs than for their master transcription factors, suggesting that tissue microenvironment is an important factor in addition to the master transcription factor in cell identity refinement. It will be interesting to investigate in future how microenvironment refines CIG expression program in a cell. We envision that one potential mechanism might be through cellular signaling involving cell-cell communications or environmental signals. In summary, SCIG is a powerful tool providing insights into cell fate determination and regulation.

## Data Availability

Our curated repository of cell identity genes along with their associated cell type annotations is accessible at https://sites.google.com/view/cigdb/curated-db?authuser=0 and Supplementary Table 1 and 2. The bulk RNA-seq datasets corresponding to the various cell types were obtained from the ENCODE project (https://www.encodeproject.org/) (29) and NCBI Sequence Read Archive (SRA) https://www.ncbi.nlm.nih.gov/sra/ (30). The accession numbers for each RNA-seq dataset are available in Supplementary Table 3. The SCIG software codes can be accessed via https://github.com/kaifuchenlab/SCIG.

## Supplementary data

Supplementary Data are available at NAR online.

## Acknowledgments

This project is supported in part by NIH grants R01GM125632, R01GM138407, R01HL148338, R01HL133254, and R01CA278832 to K.C., R01HL155632 to L.Z., and Additional Ventures (Single Ventricle Research Foundation) grant to L.Z.

## Author Contributions

K.C. conceived the project. K.C. and K.A. designed the algorithm and data analysis and interpreted the data. K.A. and B.X. wrote the codes and performed the data analysis under the supervision of K.C. and L.Z. K.A. and K.C. wrote the manuscript with comments from H.C. and L.Z. All authors reviewed and approved the manuscript.

