## Supplemental Figures for "Machine Learning Uncovers Vascular Endothelial Cell Identity Genes by Expression Regulation Features in Single Cells"

### Supplementary Figures

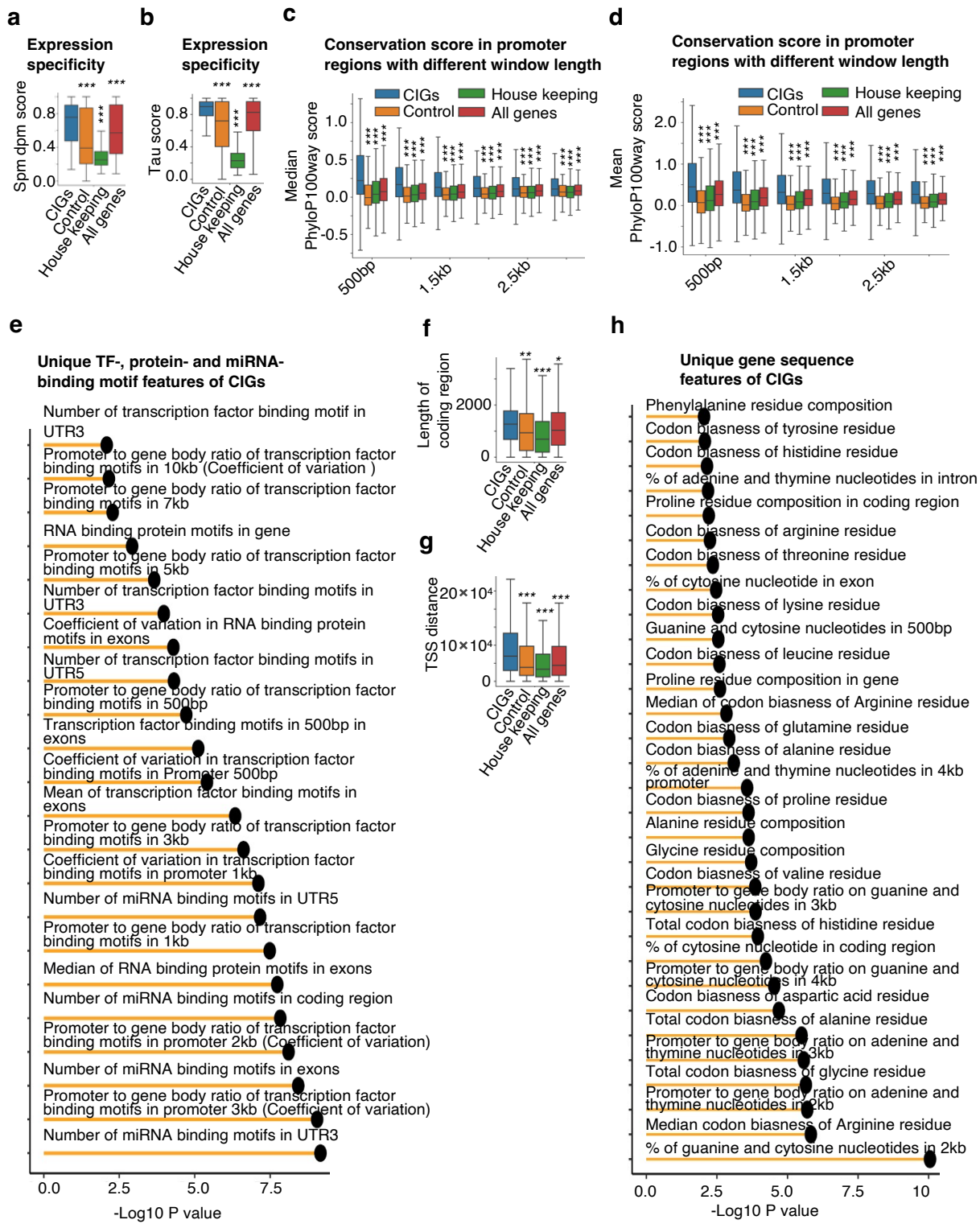

**Figure S1. Distinctive genetic sequence signatures and expression features distinguish cell identity genes from other genes. (a-b)** Box plot showing expression specificity determined by specificity measure dispersion (spm dpm) **(a)** and Tau score **(b)** for individual gene categories. **(c-d)** Median **(c)** and mean **(d)** PhyloP100way scores for promoters of individual gene categories. Scores calculated based on different promoter sizes, defined as different lengths flanking the transcription start sites, were presented. **(e)** Bar plot showing the significance of difference in individual motif features between CIGs and control genes. **(f-g)** Boxplot showing length of coding regions **(f)** and neighbor TSS distance for individual gene groups. **(h)** Bar plot showing the significance of difference in individual gene sequence features between CIGs and control genes. The two-tailed-Wilcoxon test was performed to determine the P-value of significance. \*, p-value < 0.05; \*\*, p-value < 0.01; \*\*\*, p-value < 0.001.

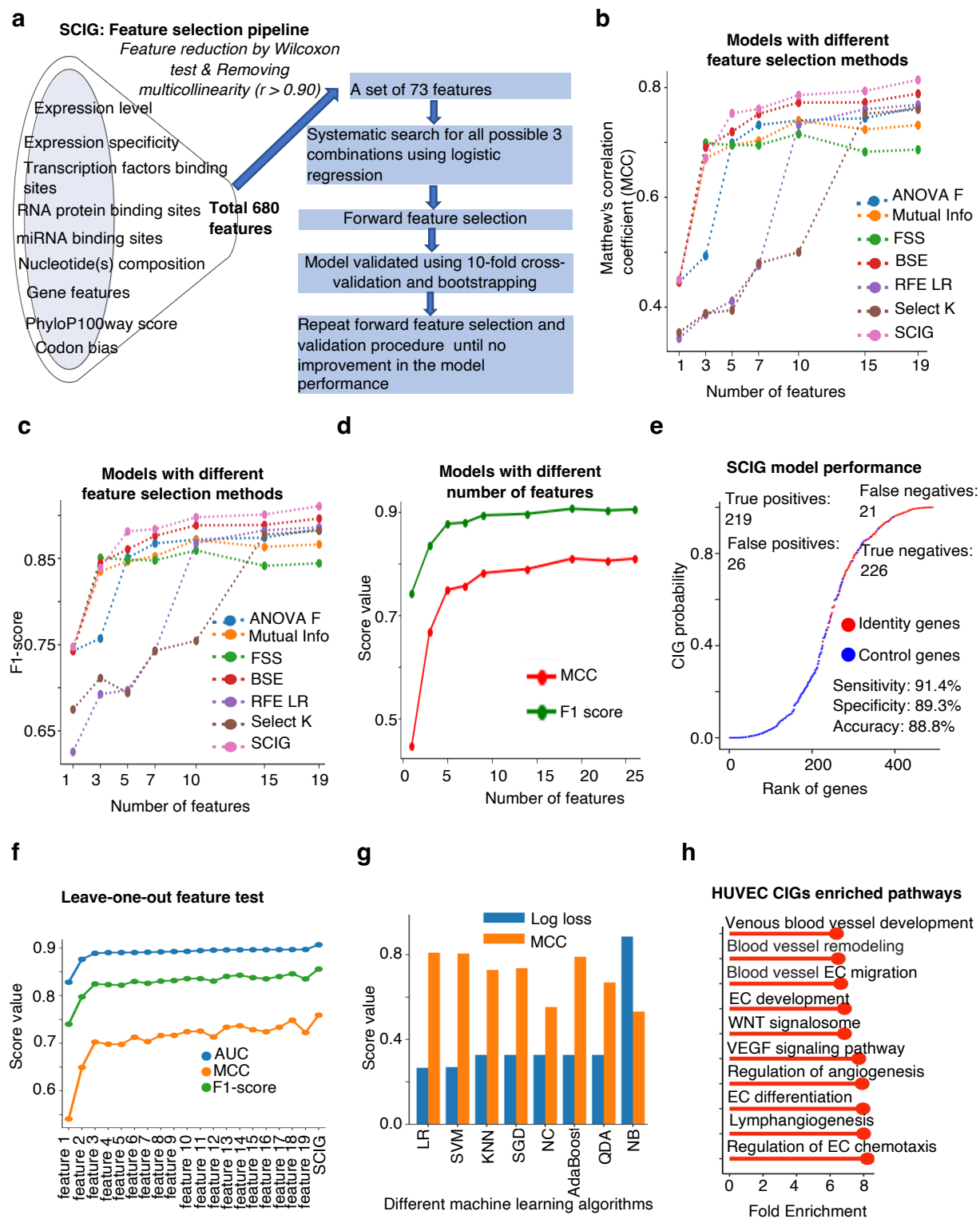

**Figure S2. SCIG is a robust machine-learning algorithm for uncovering cell identity genes.** (a) Workflow illustrating the process of feature selection in SCIG. (b-c) Line plot depicting correlation coefficient (b) and F1 score (c) as measurements of the performance of different feature selection methods, including sklearn Analysis of variance derived F-value (ANOVA F),

mutual information (MI), forward sequential selection (FSS), backward stepwise elimination (BSE), recursive feature elimination based on Logistic Regression (RFE LR), and SelectKBest (Select K). **(d)** Line plot depicting the performance of SCIG variants trained with different numbers of features. **(e)** Rank plot demonstrating the CIG probability of individual genes. **(f)** Performance of SCIG variants that were each trained with one of the 19 features depleted for comparison to the SCIG trained with all 19 features. X-axis indicates the individual depleted features. **(g)** Bar plots showing the performance of alternative machine learning algorithms to uncover CIGs based on genetic sequence signatures and gene expression features. *LR, Logistic Regression; SVM, support vector machine; KNN, K-Nearest Neighbour; SGD, Stochastic gradient descent; NC, Nearest centroid; LDA, Linear Discriminant Analysis; AdaBoost, Adaptive Boosting; NB, Naive Bayes.* **(h)** Bar plot showcasing pathway enrichment of CIGs uncovered by SCIG in HUVEC. EC, Endothelial cells; WNT, wingless-related integration site; VEGF, vascular endothelial growth factor.

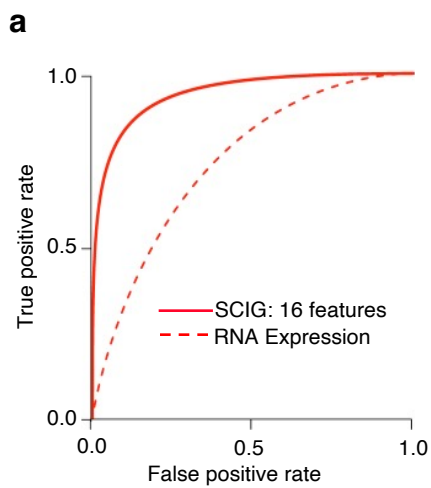

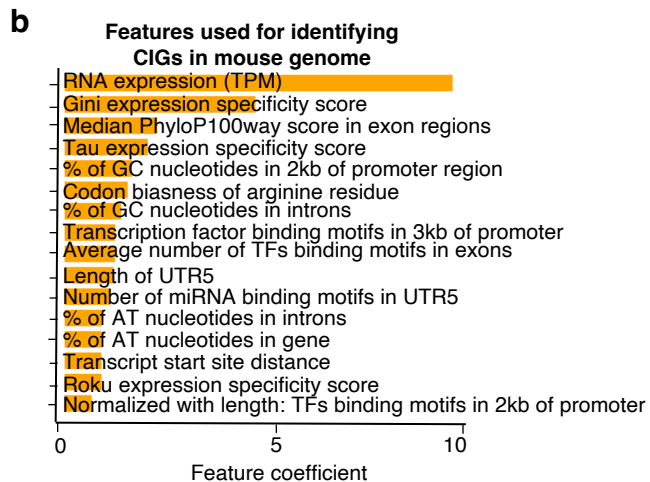

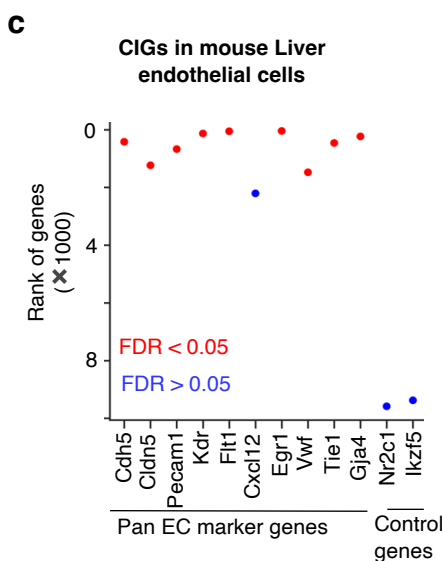

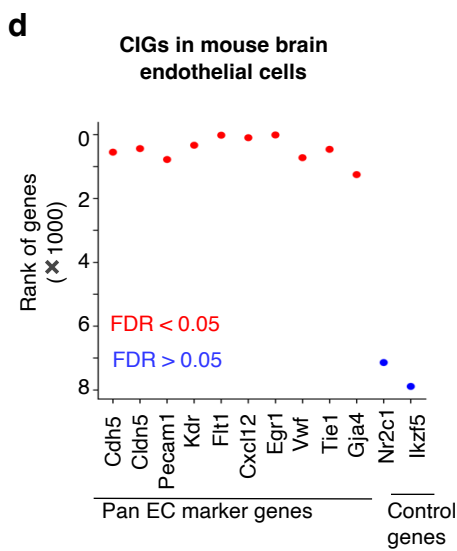

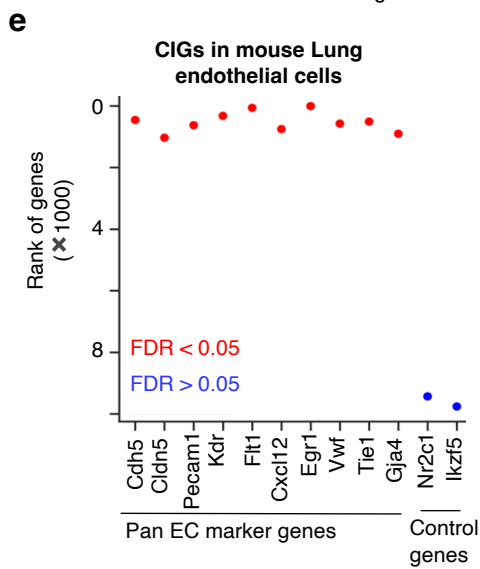

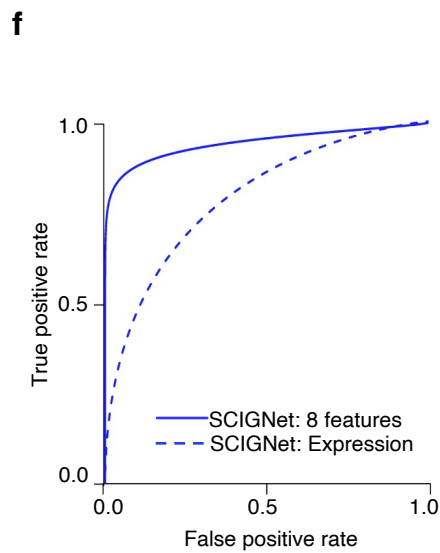

**Figure S3. SCIG uncovered cell identity genes in mouse cells with great accuracy.**

**(a)** ROC curve illustrating the performance of SCIG in mouse cells with 16 genetic sequence and RNA expression features. **(b)** Bar plot showing importance 16 significant features in the prediction of CIGs in mouse cells. **(c-e)** Endothelial marker genes ranked by CIG scores calculated by SCIG in ECs from different mouse tissue types. **(f)** ROC curve illustrating the performance of SCIGNet in uncovering the master regulators of CIGs in mouse cells.

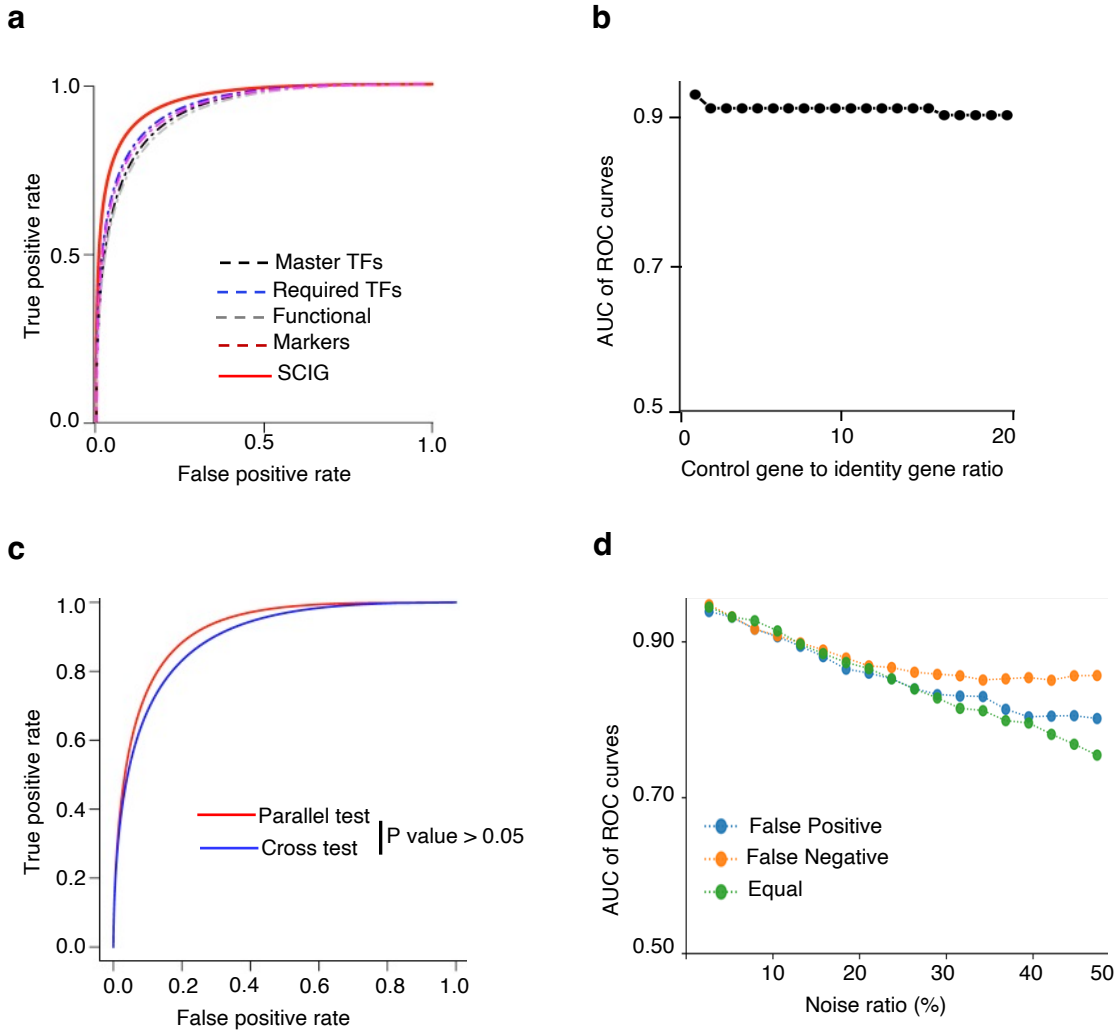

**Figure S4. Robustness and noise tolerance analysis of SCIG.**

**(a)** ROC curves demonstrating the performance of SCIG variants trained using different CIG categories but tested with all categories. **(b)** AUROC plotted against the control gene to CIGs ratio in the training data for SCIG. The ratio is systematically increased by adding more control genes while maintaining the number of CIGs constant. **(c)** ROC curves illustrating the performance of SCIG in both parallel test and cross test. In the parallel test, SCIG was trained and tested by data from the same cell types. In the cross test, SCIG was trained by data from one set of cell types and tested by data from a different set of cell types. **(d)** AUROC plotted against the simulated noise ratio in the training data for SCIG. Different methods of noise introduction are tested. These include the False Positive method, in which negative control genes are swapped to identity genes; the False Negative method, in which identity genes are swapped to negative control genes; and the Equal method, in which an equal number of identity genes are swapped to control genes and control genes are swapped to identity genes.

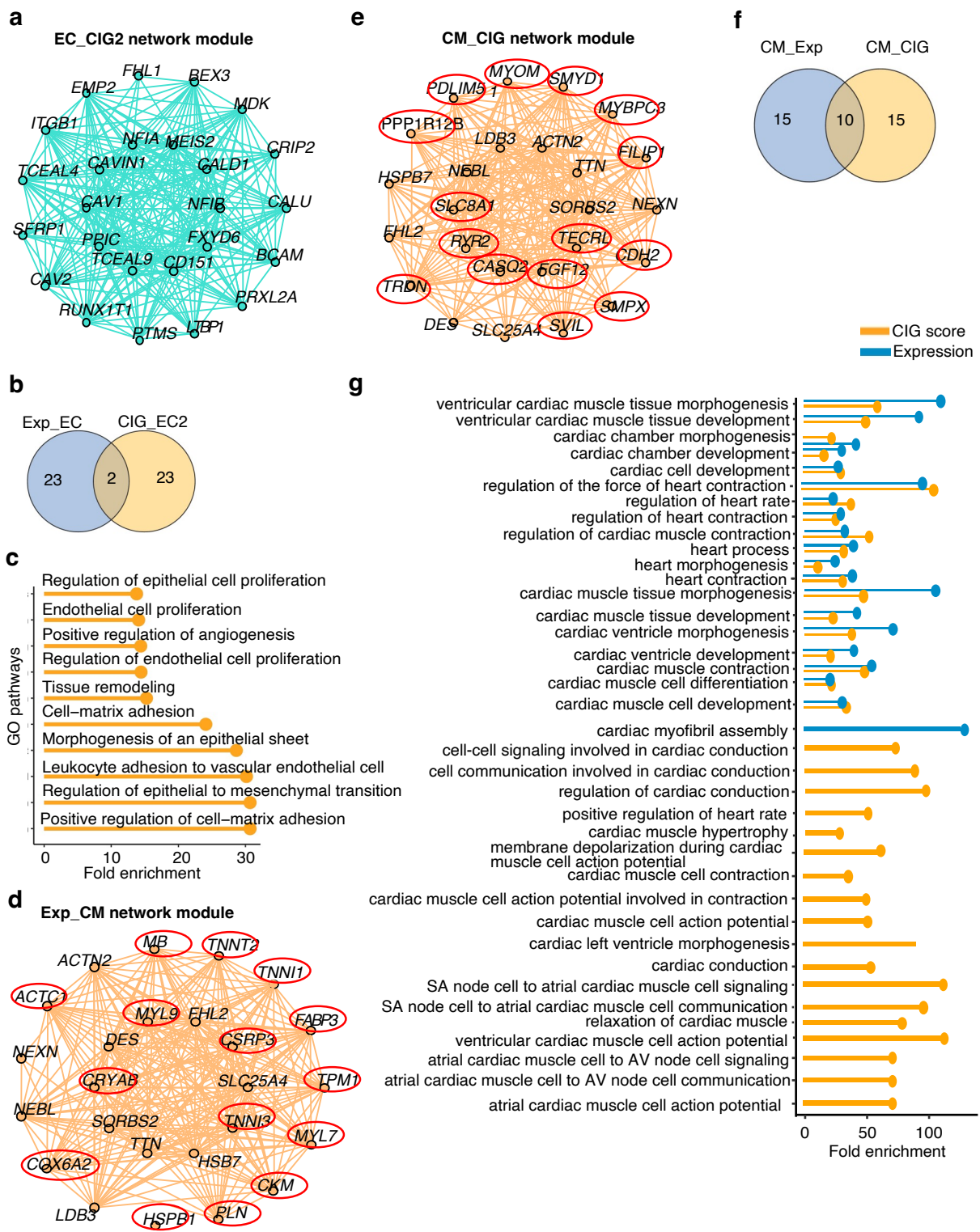

**Figure S5. Application of SCIG to gene network analysis using hdWGCNA. (a)** Network plot showing the top 25 hub genes of network module EC\_CIG2 derived from CIG score. **(b)** Ven diagram showing the overlap of hub genes in CIG score-derived network model (EC\_CIG2) and expression-derived network module (EC\_Exp). **(c)** Pathway analysis of hub genes in the CIG

score-derived network module EC\_CIG2. **(d)** Expression-derived cardiomyocytes network (CM\_Exp) hub genes. **(e)** CIG score-derived cardiomyocytes network (CM\_CIG) hub genes. **(f)** Overlap of hub genes in cardiomyocytes network modules derived from expression and CIG score. **(g)** The CIG score-derived network hub genes are significantly enriched with cardiac pathways.

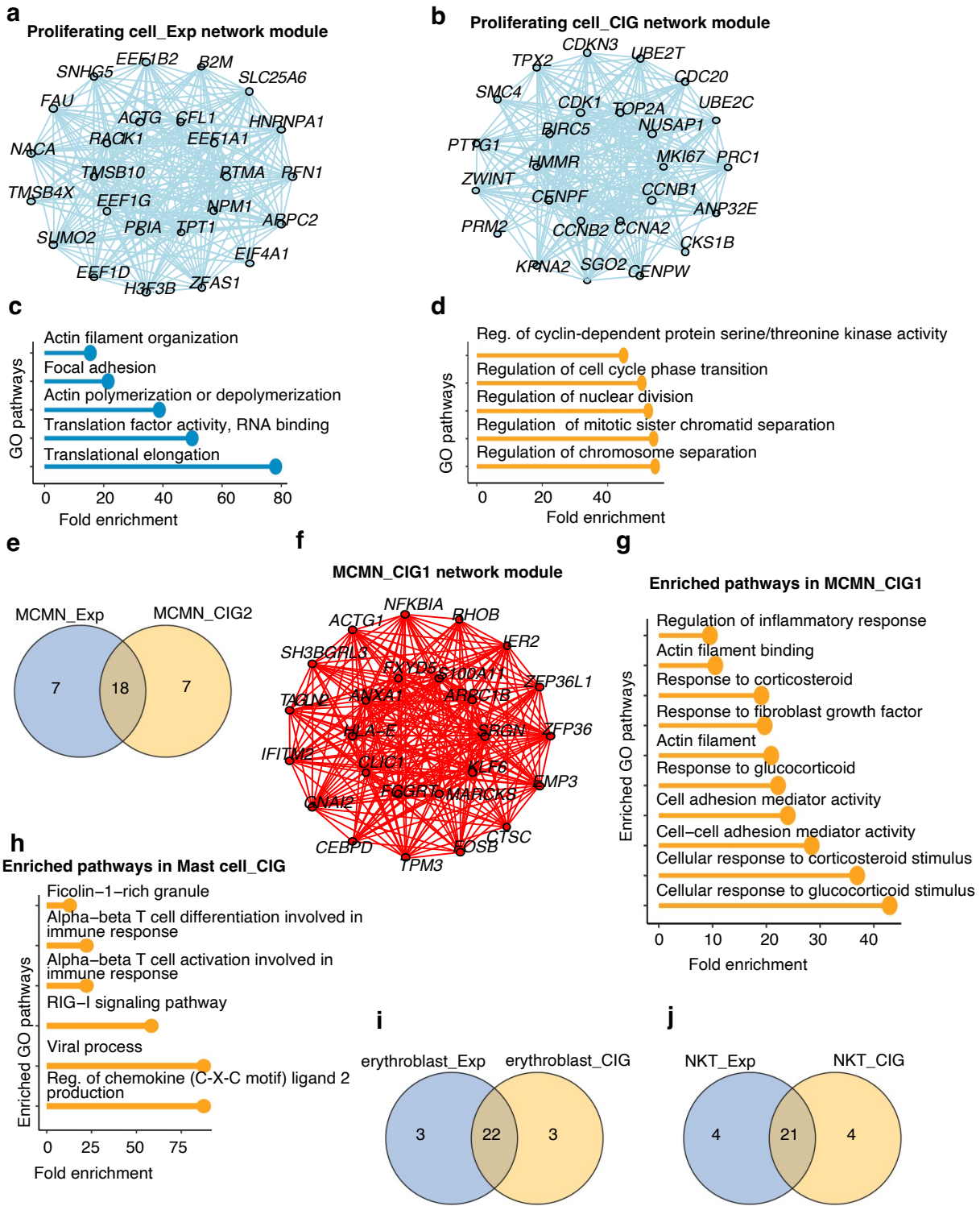

**Figure S6. CIG Score Outperforms Expression Value in Capturing Cell Identity in Network Analysis.** (a-b) Expression-derived (a) and CIG score-derived (b) gene network of proliferating cell. (c-d) Pathway analysis of hub genes from the expression-derived (c) and CIG score-derived (d) gene network of proliferating cell. (e) Venn diagram showing overlap between hub genes from the expression-derived and CIG score-derived gene network of the

macrophage/monocytes modules. **(f)** Network plot showing the hub genes of macrophages/monocytes gene network derived from the CIG score analysis. **(g-h)** Pathway analysis of hub genes in the macrophages/monocytes (g) and mast cell (h) gene networks derived from CIG score analysis. **(i-j)** Venn diagram showing overlap between hub genes between the expression- and CIG score-derived gene networks in the erythroblasts (i) and NKT (j) cells.

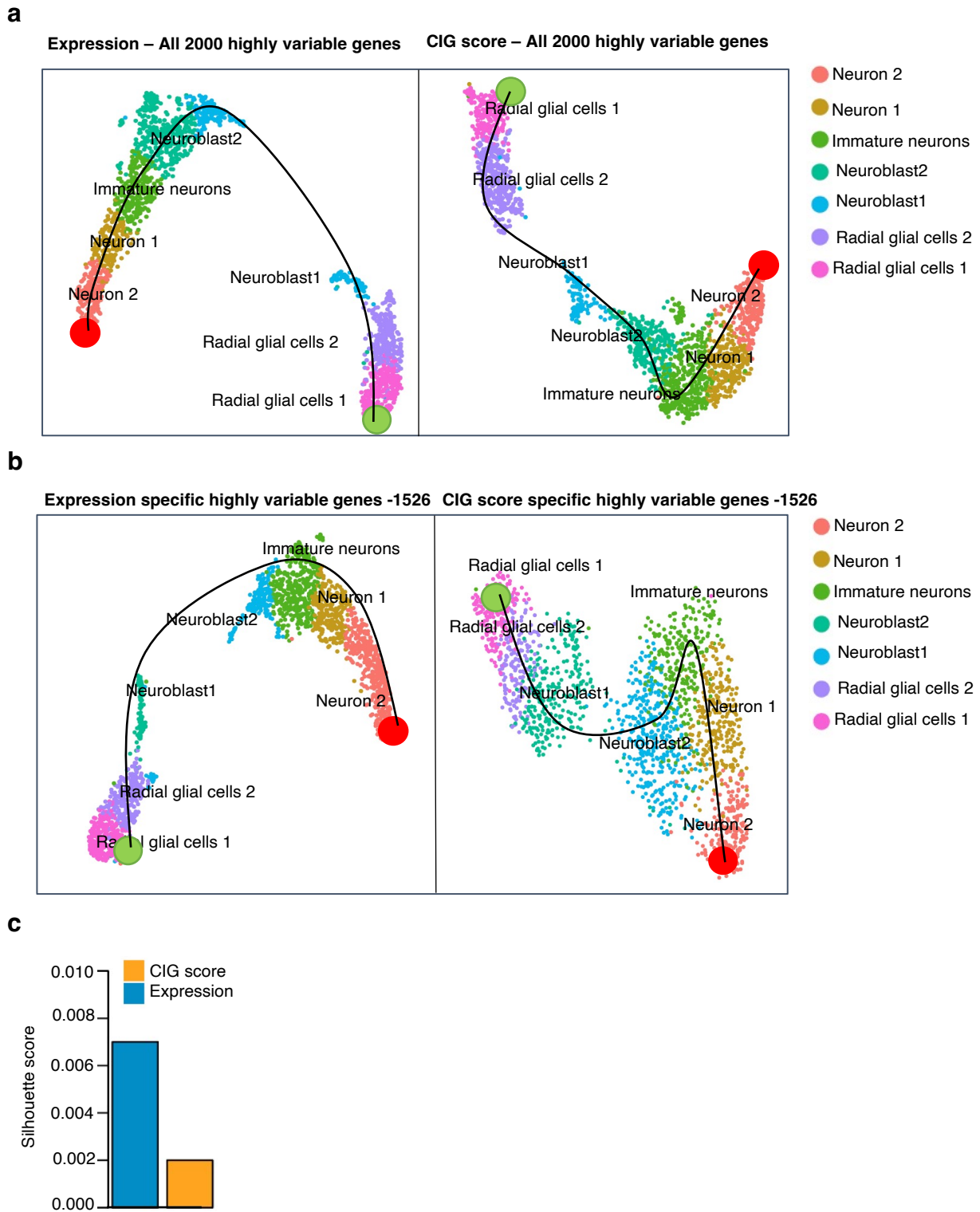

**Figure S7. Cell Identity Score Improved Single-cell Trajectory Analysis of Neuronal Differentiation. (a)** UMAP plot showing the identified cell clusters using top 2000 highly variable genes defined based on gene expression (left) or CIG scores (right). **(b)** UMAP plot showing the identified cell clusters using 1526 of the top 2000 highly variable genes defined

based on gene expression (left) or CIG scores (right) after excluding the overlapped genes between the two lists. *Green and red circles indicate locations of root and terminal cell types in trajectory analysis. The black lines indicate the differentiation trajectory.* **(c)** Silhouette score of the cell clustering based on the expression values and CIG scores.

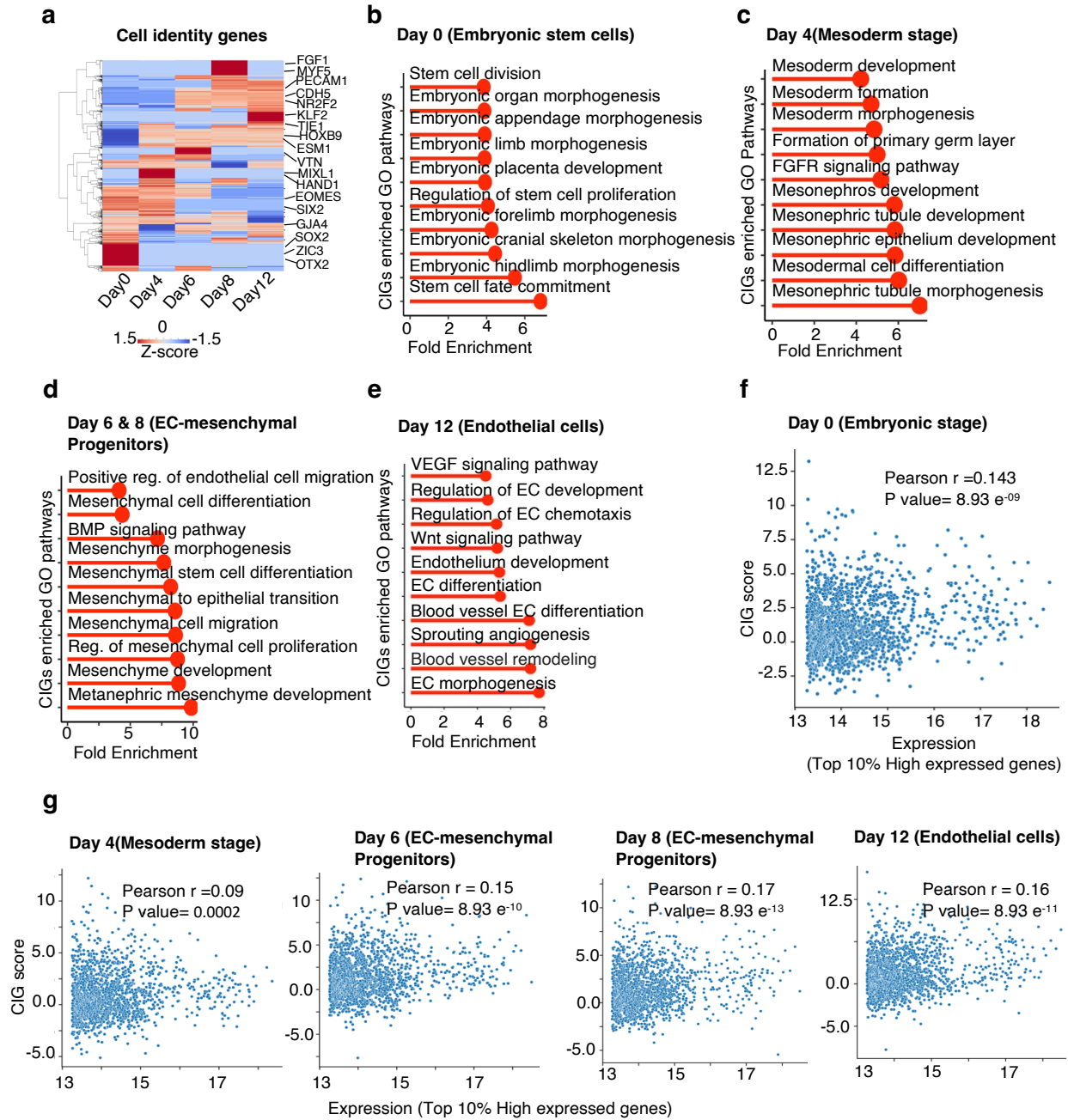

**Figure S8. Cell identity gene (CIG) landscape in the process of endothelial differentiation from embryonic stem cells (ESC).** (a) Heatmap displaying predicted cell identity genes across different stages of ESC to EC differentiation. (b-e) Pathway enrichment analysis of identified CIGs in ESC (b), Mesoderm (c), Progenitors (d), and endothelial (e) cells. (f-g) Scatter plot illustrating the correlation between expression values and predicted CIG scores of the top 10% highly expressed genes in embryonic stem cells (f) and individual differentiation stages (g).

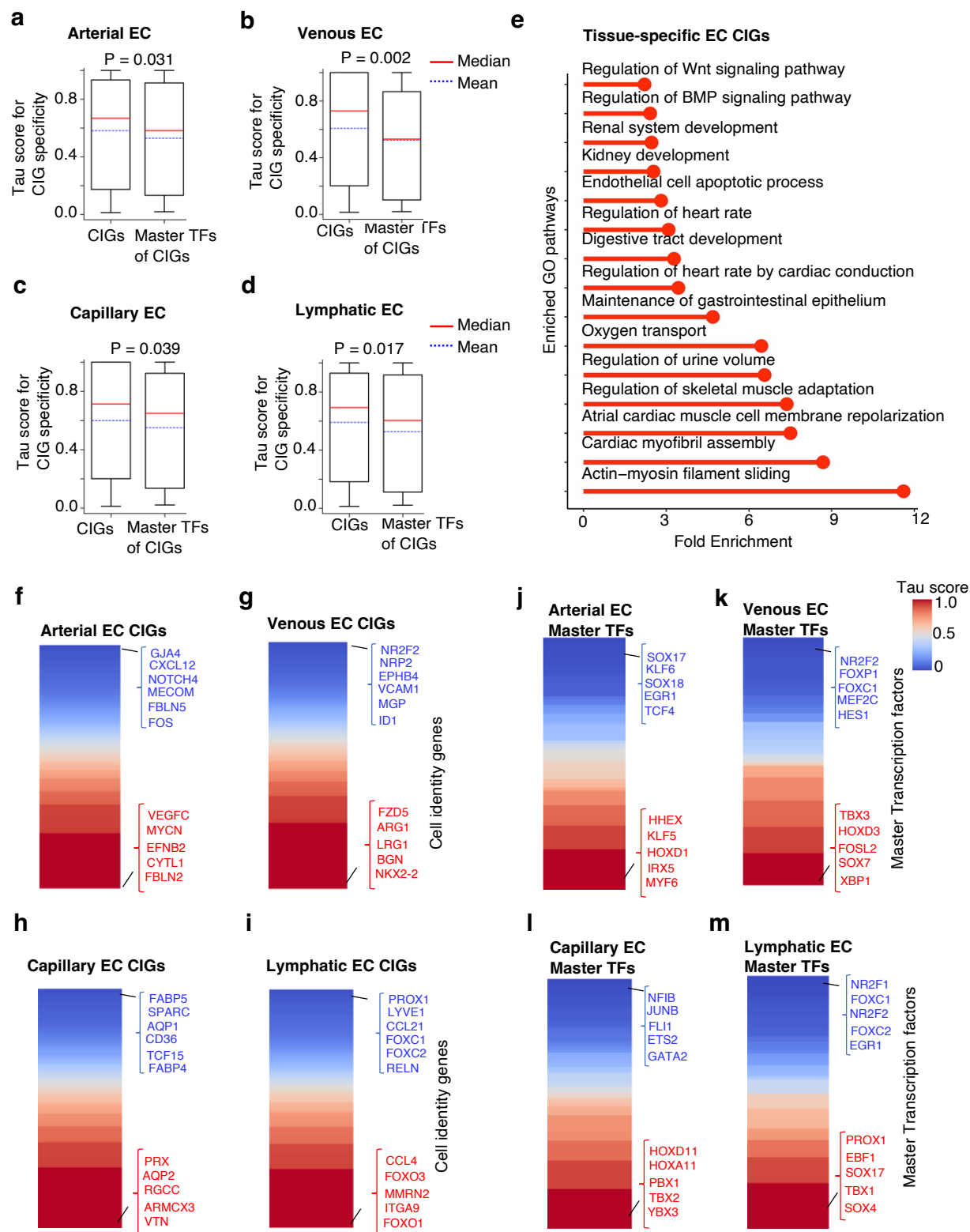

**Figure S9. Landscape analysis of arterial, capillary, venous, lymphatic endothelial cell identity genes and their master transcription factors in 15 human tissues.**
(a-d) Boxplot showing Tau score as a measurement of the expression specificity of CIGs and their regulators in individual endothelial subtypes across 15 tissues. (e) Pathway enrichment

analysis of tissue-specific endothelial CIGs. **(f-i)** Tau score computed for CIGs in individual EC subtypes across 15 tissue types. **(j-m)** Tau score computed for master transcription factors of CIGs in individual EC subtypes across 15 tissue types.
